# Collective directional migration drives the formation of heteroclonal cancer cell clusters

**DOI:** 10.1101/2022.05.18.492411

**Authors:** Miriam Palmiero, Laura Di Blasio, Valentina Monica, Barbara Peracino, Luca Primo, Alberto Puliafito

**Affiliations:** Candiolo Cancer Institute, FPO - IRCCS, Str. Prov. 142, km 3.95, 10060 Candiolo, Italy; Department of Oncology, University of Turin, 10060 Candiolo, Italy; Department of Clinical and Biological Sciences, University of Turin, San Luigi Hospital, 10043 Orbassano, Italy

## Abstract

Metastasisation occurs through the acquisition of invasive and survival capabilities that allow tumour cells to colonise distant sites. While the role of multicellular aggregates in cancer dissemination is acknowledged, the mechanisms that drive the formation of multi-clonal cell aggregates are not fully elucidated. Here we show that cancer cells of different tissue of origins can perform collective directional migration and can actively form heteroclonal aggregates in 3D, through a proliferation-independent mechanism. Coalescence of distant cell clusters is mediated by subcellular actin-rich protrusions and multicellular outgrowths that extend towards neighbouring aggregates. Coherently, perturbation of cytoskeletal dynamics impairs collective migration while myosin II activation is necessary for multicellular movements. We put forward the hypothesis that cluster attraction is mediated by secreted soluble factors consistently with the abrogation of aggregation by inhibition of PI3K/AKT/mTOR and MEK/ERK, with evidence that conditioned culture media act as chemoattractant and corroborated by a wide screening on secreted proteins. Our results present a novel collective migration model and shed light on the mechanisms of formation of heteroclonal aggregates in cancer.

## INTRODUCTION

The acquisition of invasive capabilities by cancer cells is critical to diffuse to secondary sites [1]. In tumours of epithelial origin, detachment from the primary site is frequently associated with the acquisition of mesenchymal traits [2]. A wide literature exists on the so-called epithelial-to-mesenchymal transition observed in single tumour cells, but the evidence of collective phenomena in tumour progression is also compelling. The presence of circulating tumour clusters (CTCs) in blood samples and in surgical specimens was already established in the 19^th^ century [3, 4] and recent evidence described their biological characteristics and helped elucidate their contribution to metastasis [5-9]. CTCs partially retain the epithelial traits, such as the intercellular adhesion protein, E-cadherin [10], or the expression of basal epithelial genes, as cytokeratin-14 and p63 [11]. This suggests an overlap with what happens during collective phenomena in other biological contexts such as development and regeneration [12-14].

Recent works established that metastasisation can result from the detachment of oligoclonal groups of cells from the primary tumour site other than from aggregation events in the bloodstream [5, 6]. Notably, CTCs, albeit much rarer than single circulating tumour cells, are associated with a significantly higher metastatic potential. This might be in part due to better survival rates of CTCs to anoikis and shear forces in the bloodstream [15, 16]. Moreover, heterotypic clusters of normal and tumour cells have been found in the blood circulation, and were shown to be highly proliferative and more resistant to treatments [17-19].

Survival and growth of tumour cells in both the primary and metastatic sites are associated with a number of specific alterations of the normal cell functioning, including resistance to apoptotic signals, increased migratory capabilities, paracrine interactions with stroma and stromal cells and auto-sustaining signals [2]. Cell migration away from the primary site in response to external chemical guidance cues has been reported mainly in paracrine settings where secretion is originating in the stroma and the surrounding tissues [20, 21], the blood or lymphatic circulation [22] or an organ [23, 24]. Conversely, auto-sustaining signals have been associated with both paracrine and autocrine secretion [25-29]. However, whether autocrine signalling has a role on the migratory properties of cancer cells is much less explored and remains to be fully elucidated. Recent evidence suggests that multicellular aggregates can perform directional migration [30-33] and it is proposed that gradient sensing might even be improved in multicellular aggregates with respect to single cells [34-36].

The coordinated movement of a group of cells requires cell-cell contacts retainment and the coordination of the cytoskeletal dynamics. Recent works carried out in *Drosophila melanogaster* point out the role of non-muscular myosin in collective epithelial migration, describing the importance of myosin II in the transmission of directional information and in the regulation of protrusion dynamics within and between collectively migrating cells [37, 38]. A similarly important role for myosin was also reported on mammalian epithelial cells [39].

Such data point at common underlying mechanisms in collective migration across species and biological contexts and, despite the importance of collective cell behaviour in cancer, very few simple models exist *in vitro* to explore such phenomena in detail. Growing evidence demonstrates the adequacy of 3D models to study quantitative dynamics of cell migration in various fields, such as the lymphocyte homing [40], the recruitment process of cancer cells by fibroblasts [41, 42] or cell migration of breast cancer cells [43].

Here we present a 3D model to study collective migration of cancer cells. Numerous cancer cell lines of different origins when growing in 3D show the same behaviour: they grow to form large clusters and migrate directionally toward each other through the emission of actin-rich protrusions. We called this phenotype collective directional aggregation (CDA). Quantitative measurements of the aggregation dynamics indicate that the aggregation is not due to random motion, suggesting an interaction at distance between clusters. Our results establish a crucial role for subcellular and multicellular protrusions, which are blunted by the inhibition of actin polymerisation and rely on the action of myosin contractility. We show that CDA is consistent with the action of upstream signalling, as downstream effector inhibition results in an impairment of CDA. The hypothesis that CDA is mediated by soluble attractants is confirmed by the capability of cells to migrate toward their own conditioned media and by the reciprocal attraction of heterotypic multicellular aggregates. Furthermore, a wide screening on secreted proteins confirms that conditioned media contain many chemotaxis-related factors and growth factors associated to expressed putative cognate receptors.

## RESULTS

### Cancer cell lines of different tissues of origin perform collective migration in 3D

In order to verify the universality and recurrence of CDA, we first tested a panel of 30 human cancer cell lines (listed in SI Table1) for their ability to perform active multicellular collective directional migration in 3D, i.e. multicellular aggregates merging with others. Cell lines were selected on the basis of previous knowledge about capability to grow as spheroids in a 3D matrix [44-50], which was verified with our experiments.

To this aim single cells were premixed with a hydrogel (Matrigel) in order to obtain a uniform spatial distribution. Cells and multicellular aggregates were observed by means of time-lapse microscopy for approximately three weeks after seeding. In order to monitor a large number of events and to extend our imaging experiments to a large number of samples, we performed extended depth of field (EDF) projections on wide z-stacks (details are given in the Materials and Methods section). Such technical approach was made necessary by the extreme extension of the time and spatial scales involved in the process, spanning from hours to weeks and from tens of microns to millimetres.

Representative snapshots at multiple time-points are shown in figure 1a,d (and SI Fig1A,a) and a representative movie for a subset of cell lines is reported in supplementary information (SI Movie1). Our results show that out of the 30 cell lines tested, 6 cell lines from different tissues of origin (Fig. 1a,b) are markedly different from the others in their ability to move within the matrix as multicellular clusters and display active protrusions and multicellular outgrowths (Fig. 1c). Notably, in these 6 cell lines, clusters deriving from distant single cells frequently coalesce to form “multiclonal” aggregates, forming larger and larger objects in the time span of several days (Fig. 1a from the top row to the bottom: MDA-MB-231, PC-3, ACHN, MG-63, NCI-H23 and Hs 746T). 3D collective migration assays were also performed by embedding cells in gels characterised by different protein composition, such as rat-tail type I Collagen, self-assembled synthetic peptide hydrogels and a hydrogel analogous to Matrigel. We were able to observe collective migration in matrices similar to Matrigel and in Collagen (although with differences), while cells embedded in the amorphous peptide gel proliferated within spheroids without visible signs of migration. Interestingly, while we could not detect bulk spheroids movement, multicellular aggregates seeded in Collagen were much more invasive and protrusive with respect to Matrigel, consistently with published data [51-54]. Representative snapshots and details are presented in SI Fig1B,a-c.

**Figure 1:**
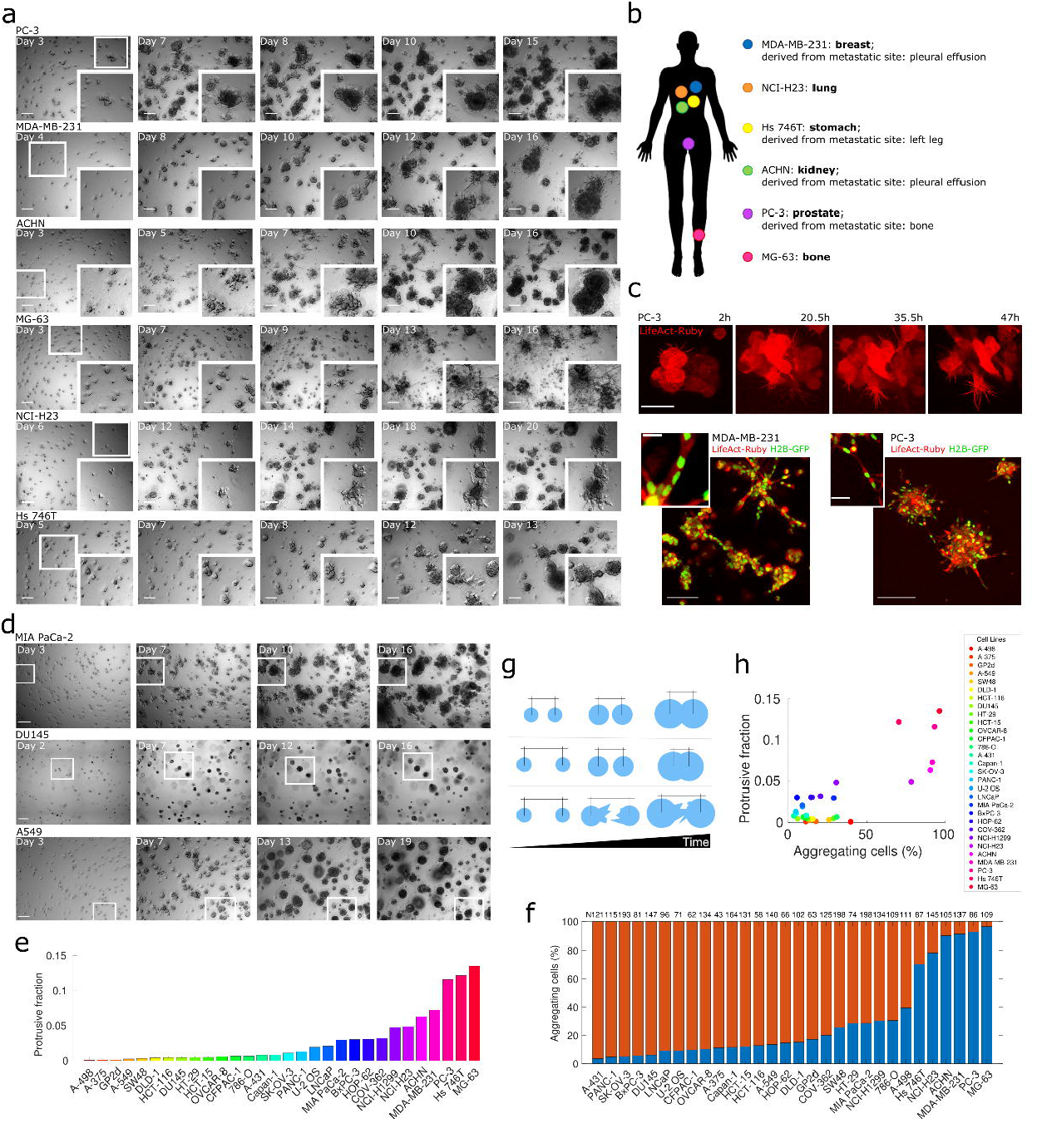
CCs grow, migrate and aggregate in 3D cultures. **(a)** Six CC lines from different tissues of origin were seeded as single-cell suspension in a 3D growth assay and imaged by means of bright-field microscopy each day for 2-3 weeks. Representative snapshots at indicated time points are shown. Each image is obtained by creating an extended depth of field (EDF) projection starting from a wide Z stack (around 1mm thick). Each row represents a cell line, from top to bottom: PC-3 (initial seeding density calculated *a posteriori*: 31 cells/mm^3^); MDA-MB-231 (22,2 cells/mm^3^); ACHN (35,1 cells/mm^3^); MG-63 (84,2 cells/mm^3^); NCI-H23 (36,2 cells/mm^3^) and Hs 746T (90,4 cells/mm^3^). The smaller white squares in the leftmost image of each row are zoomed in the corresponding inserts at the bottom right corner of the images to highlight out aggregation events. Scale bar: 200 μm. **(b)** Sketch of a human silhouette to illustrate the point of origin of each aggregating CC line. **(c)** Confocal microscopy images showing pre-aggregated 3 spheroids of PC-3 (top and bottom-right panels) and MDA-MB-231 (bottom-left panel) infected with LifeAct Ruby (top) or with H2B-GFP/LifeAct-Ruby (bottom) and seeded in Matrigel. Both subcellular protrusions (top) and long multicellular outgrowths (bottom) are well visible. Scalebar top insets: 50 μm; bottom-left inset: 100 μm; enlarge bottom-left inset: 20 μm; bottom-right inset: 200 μm; bottom-right enlarged inset: 50 μm **(d)** Representative snapshots as in panel (a) of non-aggregating cell lines. From top to bottom: MIA PaCa-2 (initial seeding density calculated *a posteriori*: 71 cells/mm^3^); DU145 (62,5 cells/mm^3^) and A459 (63,7 cells/mm^3^). White squares are meant to indicate clusters that grow without aggregating. Scale bar: 200 μm. **(e)** Ratio of protrusions to bulk regions of the spheroid for different cell lines obtained by segmentation of EDF-projected images of aggregation assays at 12 days after seeding. **(f)** Fraction of cells seeded at the beginning of the assay that are subsequently involved (blue) or not involved (red) in aggregation events over the course of an aggregation assay. “N” indicates the initial seeding densities (cells/ mm^3^) **(g)** Sketch depicting the three types of aggregation events included in the measure shown in panel (f). Light blue circles represent cell clusters, with the distance from centroid to centroid indicated with black markings on top. Top row: aggregation by growth. Middle row: aggregation by directional migration. Bottom row: aggregation by protrusive activity. **(h)** Correlation between presence of protrusions/outgrowths shown in panel (e) and the percentage of aggregating cells shown in panel (f).

In the time-lapse sequences, early after seeding, neither significant growth nor movement can be detected in our experiments. In a lag-time spanning 1 to 3 days depending on the cell line, the formation of single-and multiple-cell wide outgrowths are observed, as shown in Fig. 1c, allowing nearby initially separated objects to come into contact and move directionally one towards the other. Such directional migration events are followed by subsequent reshaping of the new “multiclonal” aggregate into a new structure, that is then compacted and assumes again the shape of a spheroidal cluster, more ordered and rounder in some cell lines (e.g., ACHN, Fig. 1a) and rich of protrusions in others (e.g., MG-63, Fig. 1a). Over the three weeks, active cluster coalescence leads to the formation of larger and larger objects, which reach the size of hundreds of microns. In contrast with the set of cell lines shown in figure 1a, other cell lines were able to grow into multicellular spheroids within the matrix, but displayed no visible and measurable signs of directional migration. In such cases coalescence of two or more clusters was only observed when objects came into contact due to growth (Fig. 1d, SI Fig. 1A,a and SI Movie2). In the vast majority of the cases, non-aggregating cell lines grow into round spheroids, presenting less or shorter outgrowths in comparison with cell lines performing CDA (Fig. 1e). It is worth noting that the phenotypic differences observed among different cell lines associated with their high or low invasiveness (i.e., the emission or long or short outgrowths and the shape more or less rounded) are consistent with what reported in literature [55-57].

In order to classify cells into aggregating or non-aggregating categories, we counted the fraction of initially seeded cells later involved in at least one event of aggregation (Fig. 1f). An aggregation event is by definition when two distinct objects (cells or spheroids) merge together forming a larger aggregate. These events can occur in three distinct prototypical ways: i) two distinct spheroids/cells grow by proliferation and the resulting increase in the radii of the spheroids drives the merging of the two objects, which do not move (Fig. 1g top row); ii) two distinct spheroids/cells actively move towards each other within the matrix and merge (Fig. 1g, middle row); iii) two spheroids emit far reaching multicellular protrusions which come into contact and drive the formation of one object (Fig. 1g bottom row).

We observed that the majority of non-aggregating cell lines only show small percentages of aggregation events (typically less than 10% at the considered densities). It is however expected that, at sufficiently high densities, all cells will present coalescence events due to the high probability of finding two objects close enough to merge due to growth. This is the reason why our aggregation assays were designed to obtain an average distance between single cells above 220 μm, thereby minimising the effect of growth-driven coalescence events. As noted above, 6 cell lines showed a significantly larger number of events (MDA-MB-231, PC-3, ACHN, MG-63, NCI-H23 and Hs 746T) ranging from about 70% (Hs 746T) to almost 100% (MG-63).

We found the fraction of aggregating cells and the emission of single cell protrusions and multiple-cell wide outgrowths to be correlated, as shown in Fig. 1h. Such correlation makes the segregation of these two groups even more evident.

Remarkably, the aggregating cell lines were originally derived from distinct cancerous tissues, as sketched in panel 1b. Moreover, different cell lines derived from the same tissue of origin (i.e., ACHN, 786-O and A-498: kidney; PC-3, DU145 and LNCaP: prostate; NCI-H23, A549, HOP-62 and NCI-H1299: lung; MG-63 and U-2 OS: bone) show different behaviours (i.e., aggregate or not), suggesting the absence of a direct correlation between the tissue of origin and the CDA phenotype.

This data suggest that this phenomenon is widespread across cancer types, independently from the tissue of origin and that it is related to phenotypic differences in the morphology of multicellular aggregates growing in a 3D matrix.

### Cluster coalescence is driven by directional collective migration

A fundamental question related to the possible biological function of collective migration is whether movement is directional or random. To this aim we developed a quantitative high-throughput approach based on the measurement of the number of isolated clusters, or cells, present in a field of view over the course of our time-lapse experiments (Fig. 2a, aggregating cells; Fig. 2b, non-aggregating cells). Each of these time series was fitted to a sigmoid function (Fig. 2c) in order to extract quantitative information on the dynamics of aggregation in each assay separately.

**Figure 2:**
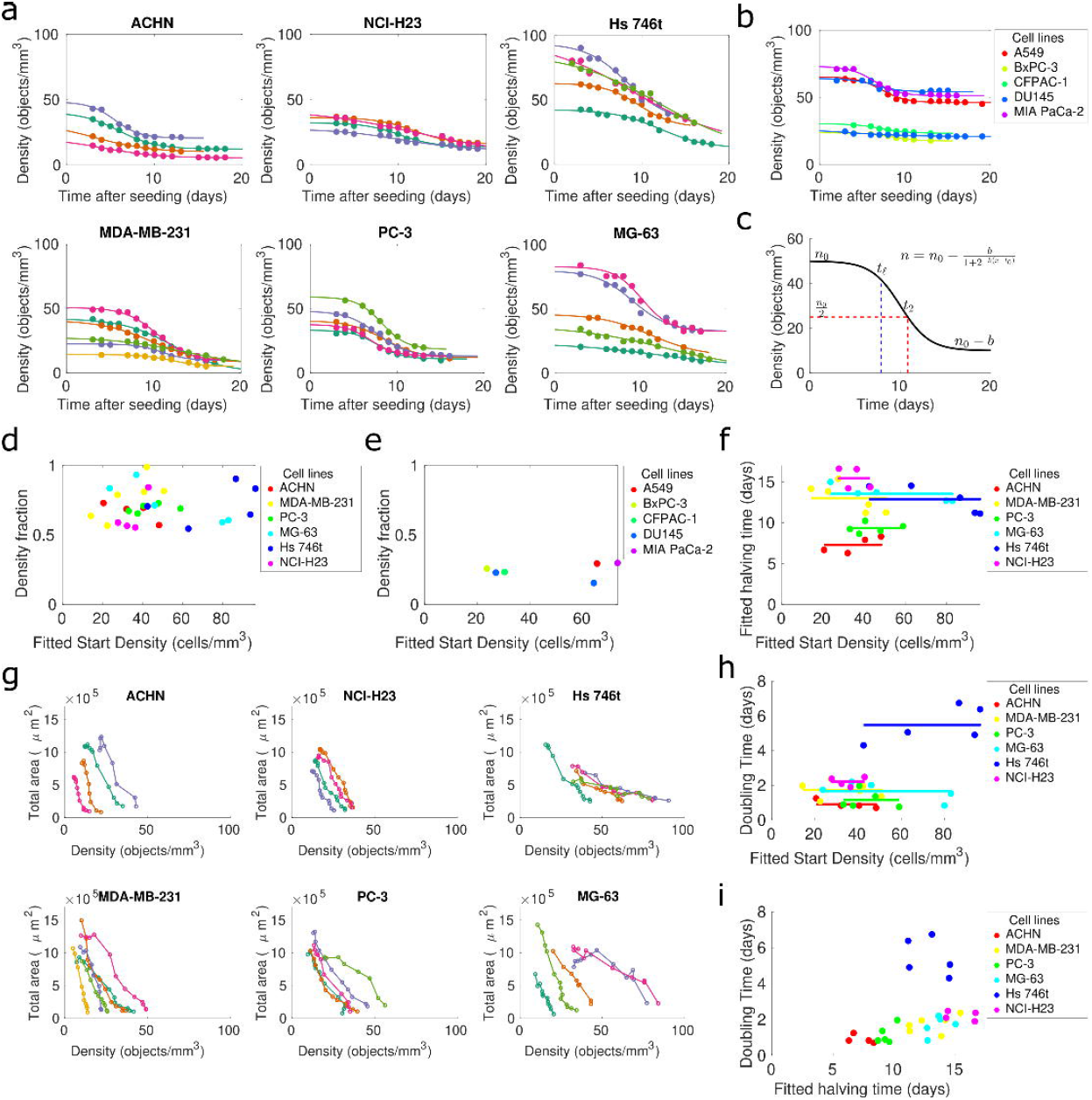
Aggregation kinetics is independent on initial seeding density and coupled to cell proliferation rates. **(a)** The number of separate objects in each EDF-projected image over time is fitted with a sigmoidal curve for different initial seeding density. Each set of lines and points represent a single aggregation assay and points are experimental data, while lines represent the fit. **(b)** Same of (a) but for non-aggregating cell lines. The legend indicates the colours associated to cell lines. **(c)** Plot of the sigmoid function used to fit the data of aggregation kinetics with indicated parameters used in the quantification. The operational definitions of each parameter are: halving time is the time at which the number of objects is half the initial fitted number of objects; lag time t_l is the time corresponding to the first change of convexity of the function; n_0-b is the number of objects at the end of the assay. **(d**,**e)** The density fraction (difference between number of objects at the beginning and the end of the assay as resulting from the fit parameters divided by the number of objects at the beginning) is plotted for each cell line and initial starting density. **(f)** The plot reports the halving times obtained by the fit parameters of curves shown in panel (a) and plotted as a function of the seeding density. **(g)** Scatter plots reporting fitted starting density vs the total area of the growing spheroids, i.e., the total projected area in an EDF-projected field of view. Each dot represents a single time-point from the aggregation assays. Lines parallel to the y axis would indicate pure proliferation with no aggregation, while lines parallel to the x-axis would indicate aggregation with no proliferation. **(h**,**i)** The plots report the approximated doubling times (inferred from time-series of total projected area), and the scatterplot halving time vs doubling time obtained by the fit parameters of curves shown in panel (a) and plotted as a function of the seeding density. Doubling times were obtained by a separate fit (see details in the Materials and Methods section, and growth curves reported in SI 2a). Each dot represents an aggregation assay performed with a cell line at a given initial seeding density with colour codes reported in the legend.

We first ought to quantify the extent of the aggregation phenomenon in different cell lines and for different initial seeding densities. This measure is given by the ratio between the variation of the number of separated objects at seeding with respect to the end of the assay, and the seeding density. Such a measure, which was termed density fraction, would be vanishing for non-aggregating cells, while it would display values closer to unity for strongly aggregating cells. Density fractions for a set of cell lines are shown in panels 2d and 2e. As expected, aggregating cell lines display a decrease in density which correspond to final densities smaller than half of the initial density (Fig. 2d). In contrast, non-aggregating cell lines do not show significant decrease in density, as visible also from the curves in Fig. 2b and in Fig. 2e. As explained above, a small number of coalescence events can be seen even in non-aggregating cell lines due to growth and random positioning of the cells at seeding, as shown by the small density fractions plotted in Fig. 2e (see also sketch in Fig 1g).

To verify whether aggregation might be induced by random movement, we measured the effect of the initial distance between cells at seeding (i.e., density) on the dynamics of the aggregation process. To this aim we developed a way to quantify the kinetics of the aggregation process by measuring the halving time, i.e., the time it takes to halve the number of distinct objects in a given volume with respect to the number at seeding. We have shown previously [32] that random or directional movement would have very different impacts on the kinetics of aggregation. If the movement of clusters (or the directionality of protrusions and outgrowths) were random, higher densities would lead to a larger number of aggregation events simply due to the fact that, when cells sit closer, the chance of coming into contact is higher. In particular, this would imply an inverse proportionality between the halving times and the seeding density [32].

We therefore measured the halving times of cell lines at different initial seeding densities, as shown in Fig. 2f. Our results show that the aggregation halving times are largely independent of initial seeding density, therefore indicating that migration is not random. Aggregation times show variability across different cell lines and fitted data indicate that ACHN and PC-3 are the fastest to aggregate, with halving times of (7.3±1.0) days and (9.4±0.7) days respectively (mean±std); MG-63, Hs 746T and MDA-MB-231 halve in (13.6±1.0) days; (12.9±1.7) days and (13.0±1.7) days respectively and NCI-H23 is the slowest in the group, with a halving time of (15.5±1.3) days. Overall, this phenomenon occurs on a timescale of the order of a week for all cell lines considered. This is consistent with what previously measured for PC-3 cells [32], and is also significantly different from what observed in liquid overlay cultures, where random movement is predominant [58-60], and the dynamics is much faster, with aggregation times estimated as less than a day.

### Aggregation is coupled with proliferation of cells within clusters

To dissect whether proliferation and aggregation occur on the same timescale, we quantified the growth of clusters during the aggregation assays (SI Fig. 2a,b) as inferred by the total projected areas in the time-lapse experiments, and plotted it against the kinetics of aggregation, as shown in Fig. 2g and SI Fig. 2c for non-aggregating cell lines. These scatterplots show clearly that separating the effect of proliferation from that of aggregation is a daunting task in aggregation assays starting from monodisperse single-cell size distributions, as these two phenomena occur on similar timescales. Indeed, if there was separation of scales, the scatterplots in Fig. 2g would show points all aligned with one or the other axis. On the contrary, growth and aggregation occur simultaneously as witnessed by sequences of points that do not lie parallel to the coordinate axes. Therefore, in order to clarify this relationship, and to investigate potential correlations between proliferation and aggregation, we measured the approximated doubling time of spheroids (calculated from their projected area) as shown in Fig. 2h and SI Fig. 2d for non-aggregating cell lines. Reported means±std for each cell line are ACHN (0.91±0.24) days; MDA-MB-231 (1.74±0.47) days; PC-3 (1.16±0.51) days; MG-63 (1.67±0.53) days; Hs 746t (5.48±1.04) days and NCI-H23 (2.21±0.26) days. Approximated doubling times for aggregating cell lines are consistent with proliferation rates on flat cultures (1.25 days (ACHN); 1.25 days (MDA-MB-231); 1 day (PC-3); 1.25 days (MG-63); 1.6 days (NCI-H23) [61] and 2.8 days for Hs 746T [62]). Furthermore, coherently with our hypothesis, halving times and doubling times correlate, and faster aggregating cells are found to be on average also more proliferating in our 3D single cell assay (Fig. 2i). These observations hold for all cell lines but for Hs 746T, which show halving times comparable to other cell lines but much slower doubling times. Correlation coefficients for halving and doubling times are indeed R=0.345, p=0.067 for the complete dataset, and R=0.752, p=2.25×10^−5^ excluding Hs 746T. These observations are coherent with the notion that it is indeed hard to disentangle the effect of proliferation from that of directional migration in this experimental setup which spans several doubling times for all cell lines.

In order to further dissect the relationship between proliferation and aggregation, we developed an aggregation assay starting from pre-formed multicellular aggregates, to decrease the relative importance of proliferation. Cell spheroids were pre-assembled under non-adherent culture conditions and were embedded in the same hydrogel as for single cell assays.

With this experimental setup, the emission of protrusions and collective migration start not later than 24h after seeding. The overall estimated change in the total number of cells over the course of single cell assays was around 10-fold in all cell lines, while spheroid assays involved at most 2-fold changes. Therefore, consistently with the brevity of the assay and the larger structures involved, growth of spheroids during the assay was more limited than in single cell assays. These observations are therefore consistent with the notion that, in pre-assembled spheroids assays, directional migration is occurring on faster scales than growth. Our experiments show that spheroids seeded in a 3D hydrogel start emitting single cell protrusions and multicellular outgrowths already a few hours after seeding and migrate collectively forming multi-cluster aggregates as shown in Fig. 3a and in SI Movie3. All aggregating cell lines recapitulate the same behaviour observed in single cell assays, while non-aggregating cells only show growth, with no signs of collective migration (Fig. 3b).

**Figure 3:**
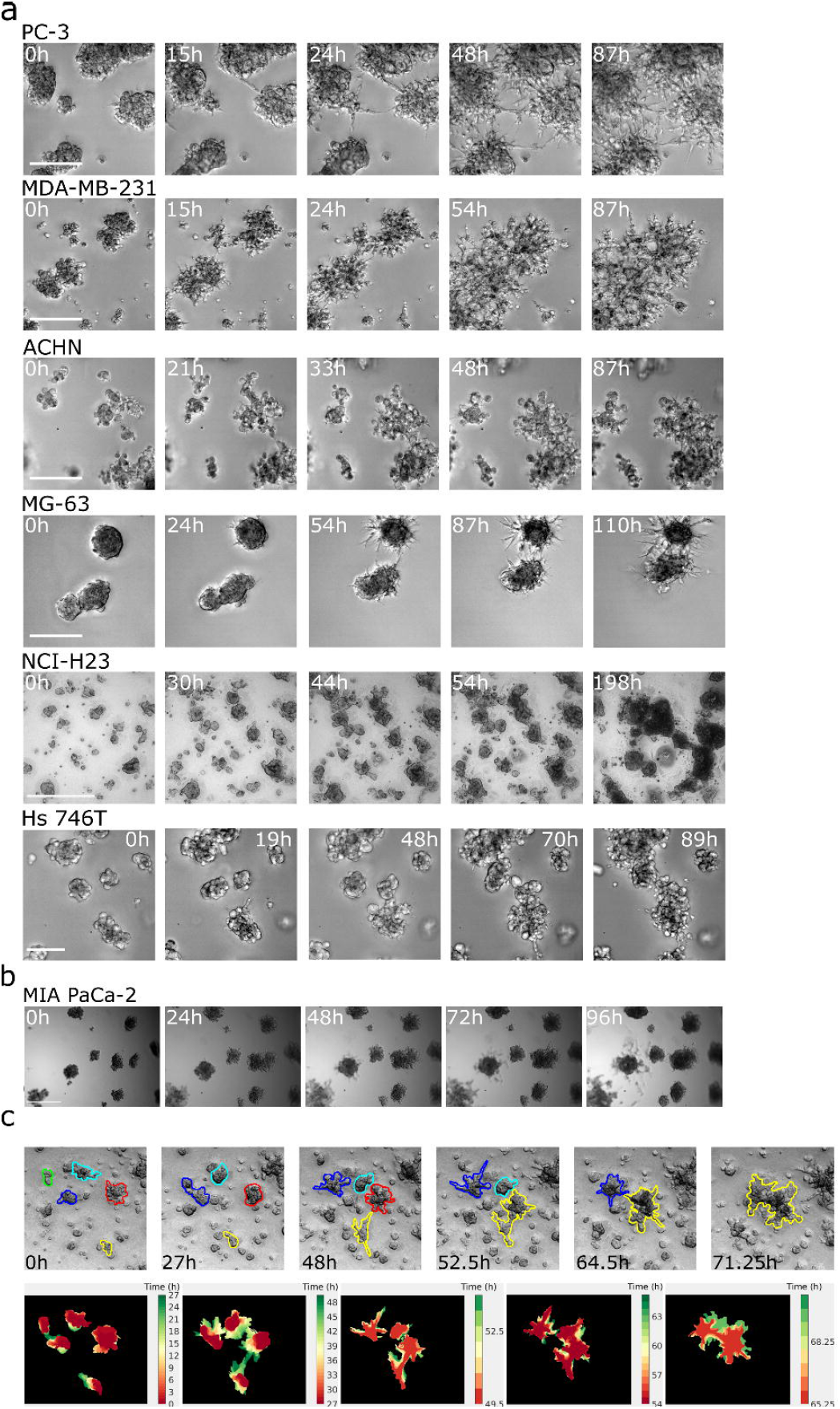
Pre-formed cell spheroids perform CDA. **(a)** Each row shows representative snapshots of a time-lapse experiment with an aggregation assay performed starting from pre-aggregate spheroids (the same six cell lines showed in Fig. 1a are shown). Seeding density: 2,5 spheroids/mm^3^. Scale bar: 500 μm. **(b)** The pre-formed spheroids aggregation assay performed with the MIA PaCa-2 cell line (among the non-aggregating cells). A single merging event is observed when two spheroids come into contact due to growth. Seeding density: 2,5 spheroids/mm^3^. Scale bar: 500 μm. **(c)** Top: representative snapshots of an aggregation assay with pre-formed spheroids where 5 clusters were manually segmented and tracked over the course of the assay. Each colour defines the outline of an object, and one of the two colours of two merging objects is retained in an aggregation event. Bottom: the outlines of the spheroids (sampled every 3 hours) as in the top row were superposed with a time dependent colour code (red: earliest time-point; green: latest time-point) in order to highlight the protrusions mediating the aggregation event. Each panel ends when two spheroids merge.

Despite starting from different initial conditions, the two assays show the same collective migration paradigm, with single and multicellular protrusion mediating the coalescence events as shown in the insets of Fig 3a.

To pinpoint the directionality of the movement, we segmented a group of aggregating objects in the images of our time-lapse assay for the entire duration of the aggregation assay and extracted the shape of the projected area occupied by the objects over time (Fig. 3c). Our results show that spheroids move directionally toward each other to form one single cluster at the end of the movie and that spheroid deformation and protrusions occur along the line separating the two objects, albeit with considerable fluctuations in the orientation. This observation is consistent with what observed in other biological settings [63], where protrusions are not statically oriented towards the source, but are dynamically moving, maintaining overall orientation in time.

### Collective directional migration is not driven by active cell proliferation

To further clarify the effect of growth on collective directional migration we performed a series of experiments to decouple proliferation and migration. To this aim we treated cells with mitomycin C, a chemotherapeutic drug with known ability to block cell cycle progression. In order to avoid undesired effects of the drug, a number of controls were performed. First, we ought to determine effective concentrations and treatment duration to induce effective proliferation block with no or minimal effect on cell viability. To this aim we treated cells with mitomycin at concentrations ranging from 0.3 μg/ml to 2 μg/ml and assessed growth (or lack of) by monitoring the number of adhering cells in a time-lapse experiment, as shown in Fig 4a. The number of adhering cells was fitted with an exponential to derive a net growth rate, i.e. the difference between cell doubling rate and cell death rate. Simultaneously, a live apoptosis detection dye was used to separately evaluate toxicity of the treatment, as shown in SI Fig. 4a.

**Figure 4:**
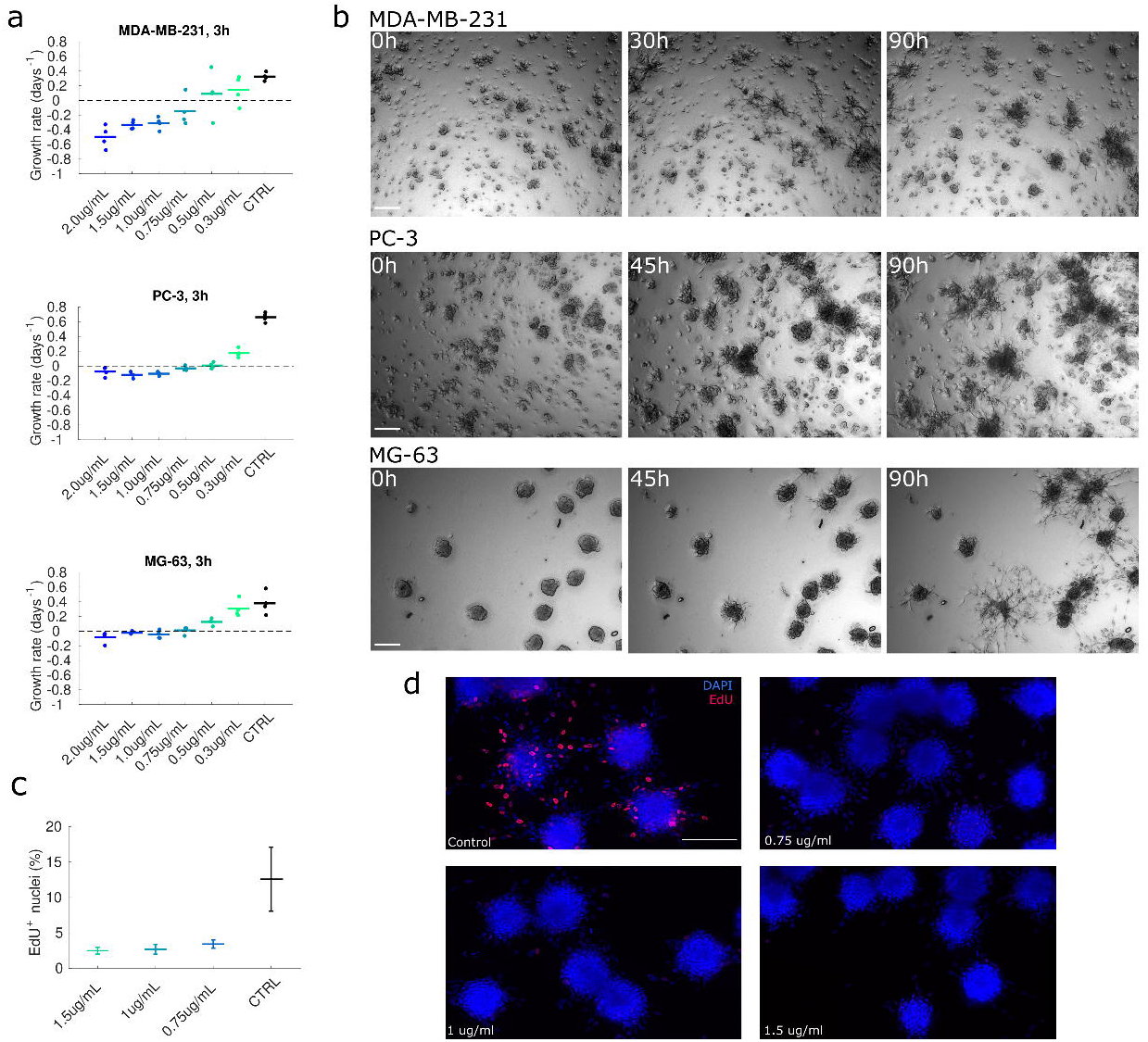
CDA proceeds independently from proliferation. **(a)** The plots show the growth rate fitted by measuring the number of cells by image segmentation in cell lines cultured in 2D and treated with mitomycin at the indicated concentration. Positive growth rates indicate the presence of cell proliferation while negative growth rates indicate toxicity. The duration of the time-lapse is 19 hours. Note that growth rates in control conditions are consistent with doubling times measured in aggregation assays. **(b)** Snapshots of spheroid-based aggregation assays performed with mitomycin-treated cells (0.75 μg/ml). Scale bar 200 μm. **(c)** Fraction of EdU incorporating cells in deconvolved images of spheroids treated with mitomycin for MG-63 spheroids as those shown in panel (d). Each point is the mean of the fractions obtained by image segmentation in 4 different fields of view. **(d)** Snapshots of aggregating spheroids stained with EdU and DAPI for MG-63. Scale bar 200 μm. Images are equalised with the same limits.

This set of experiments helped us developing the treatment conditions to perform the spheroid-based aggregation assay with cells treated with mitomycin. Spheroids were treated for 3h with mitomycin right after seeding and then put under a microscope for time-lapse observations. Representative snapshots of spheroids treated with mitomycin are shown in Fig. 4b and full time-lapse movies are included as SI Movie 4-6. The effect of mitomycin on proliferation in the 3D spheroid-based assay is shown by a staining with EdU (Fig. 4c,d), which indicates drastically reduced proliferation in treated spheroids. In conclusion, blunting cell proliferation has no apparent effect on collective directional aggregation, confirming that while these two phenomena are coupled when both present, they are indeed not causally related. In particular, our experiments show that proliferation and growth do not drive collective directional aggregation.

### The molecular perturbation of cytoskeletal components impairs the aggregation process

To corroborate the observations on the important role of subcellular and multicellular protrusions reported and discussed in the previous sections, we performed higher resolution microscopy experiments with more specific molecular markers. To this aim we stably expressed LifeAct-GFP or LifeAct-Ruby in a subset of cell lines, and observed single aggregation events. Under these experimental conditions, the actin content of both single and multiple cells outgrowths was well visible in the form of actin filaments.

Cells expressing either LifeAct-GFP or LifeAct-Ruby were seeded as pre-formed spheroids in Matrigel and imaged by means of fluorescence microscopy for several days in order to distinguish the separate contributions of each cluster. Representative snapshots reported in Fig. 5a (see also SI Movie7) show that the aggregation between two neighbouring, initially separated, objects is accompanied by the reciprocal emission of actin-rich protrusions oriented along the longitudinal axis of the two spheroids.

**Figure 5:**
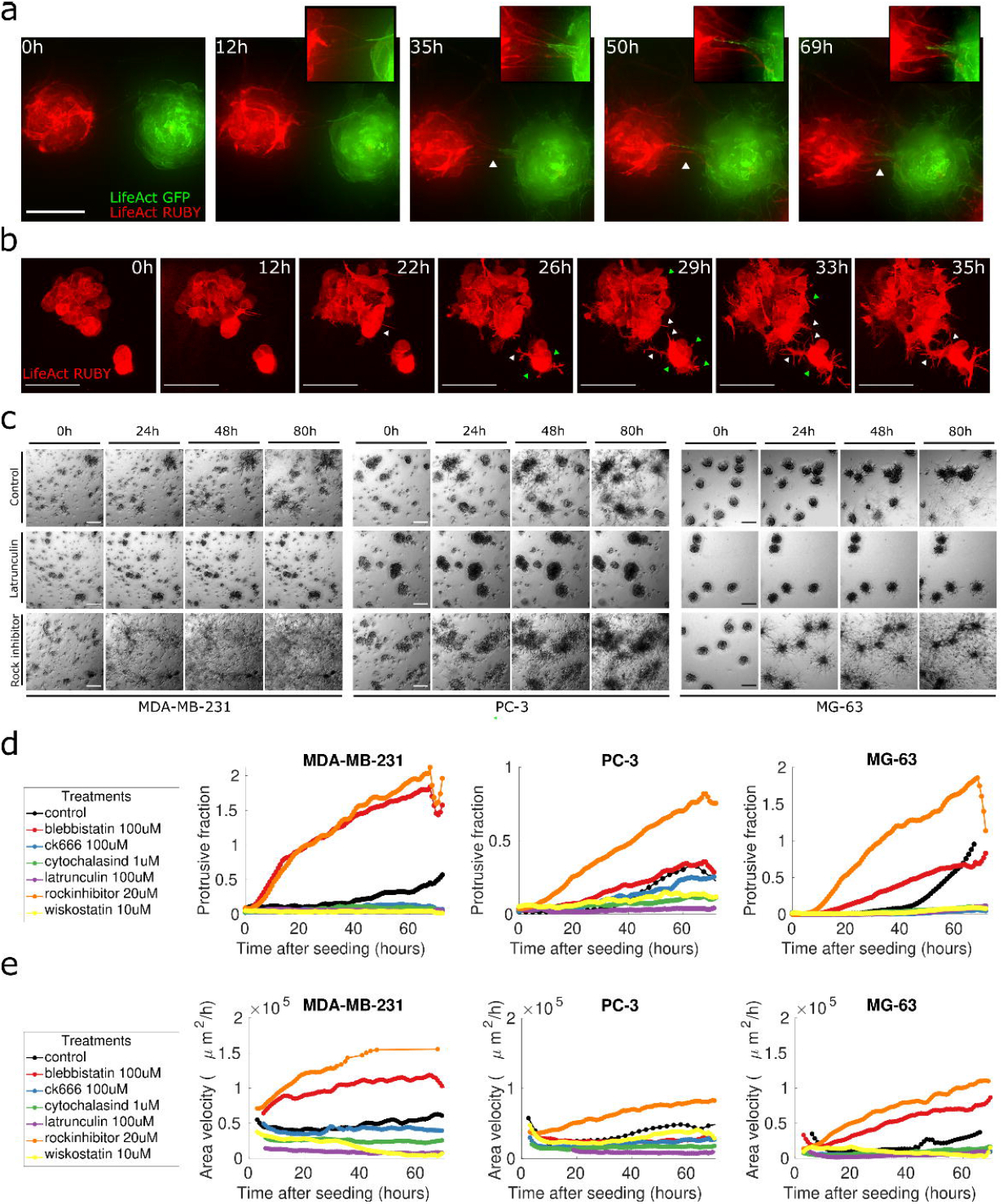
The role of protrusions in CDA. **(a)** Representative images showing pre-formed spheroids of MG-63 cells transduced with LifeAct-GFP (green) or LifeAct-Ruby (red) expressing lentiviral vectors, embedded in Matrigel. White triangles indicate protrusions between the two spheroids. The region of the images corresponding to protrusions was enlarged and shown in the top-right inserts. Scale bar: 100 μm. **(b)** Representative images of pre-formed spheroids of LifeAct-Ruby expressing MDA-MB-231 embedded in Matrigel. White triangles point at protrusions directed toward the neighbouring object. Green triangles point at smaller protrusions which develop all around the spheroids. Scale bar: 100 μm. **(c)** Pre-formed spheroids of three selected aggregating cell lines were either left untreated or treated with 1 μM Latrunculin A or 20 μM Y-27632. (From left to right: MDA-MB-231, PC-3 and MG-63; from top to bottom: untreated, Latrunculin A, Y-27632). Snapshots are extracted from time-lapse experiments at 0, 24, 48, and 80 hours after seeding. Seeding density: 2,5 spheroids/mm^3^. Scale bar: 200 μm. **(d)** The images corresponding to experiments as those shown in panel (c) were segmented in order to extract the ratio of protrusion to bulk regions for each treatment, as detailed in the legend. **(e)** Area velocity extracted from images as detailed in the Material and Methods section. Each colour represents a different treatment as detailed in the legend.

Confocal microscopy time-lapse experiments (Fig. 5b and SI Movie8) allowed us to investigate in detail the morphological characteristics and the dynamics of the protrusions. We observed that spheroids of cell lines able to perform CDA, emit actin-rich long protrusions, which are already visible only a few hours after seeding. Small subcellular protrusions are emitted all around the spheroids (green triangles in Fig. 5b), while longer, several cells-wide, outgrowths develop in the direction of the neighbouring spheroid (white triangles in Fig. 5b) and allow the two different objects to enter in contact and to subsequently merge.

To further confirm the importance of protrusions for directional migration to occur, and to investigate the role of the cytoskeletal machinery in the aggregation process, we tested whether we could perturb CDA by inhibiting crucial cytoskeletal components.

To demonstrate that actin polymerisation is required in CDA, we used four inhibitors with different mechanisms of action: Latrunculin A and Cytochalasin D as actin polymerisation inhibitors; CK-666 as an actin assembly/branching and Wiskostatin as a WASP inhibitor, perturbing actin polymerisation. Likewise, to test the role of myosin II, we used Blebbistatin which is an inhibitor of myosin-II ATPase activity and the ROCK inhibitor Y-27632, which prevents the phosphorylation of myosin II light chain (MLC) mediated by ROCK, a major downstream effector of the small GTPase RhoA.

To perform the experiments, a selected set of cell lines were tested in pre-formed spheroids assays. This choice was necessary to reduce the duration of the assay and to minimise the effects of the drugs on other important cellular processes. The images of the treated spheroids are reported in Fig. 5c and SI Fig. 5a and show that in all the selected cell lines, both actin and myosin perturbation lead to the impairment of CDA (see also SI Movie9-15). In order to appreciate the effect of all these treatments we quantified both the emission of protrusions (Fig. 5d) and the presence of movement (Fig. 5e) by implementing image analysis algorithms based on digital segmentation and tracking.

The effects of actin perturbation on CDA can be appreciated from the representative snapshots of time-lapse experiments (Fig. 5c, second row and SI Fig. 5a), where all the treatments blunt the protrusive activity and the capability of the spheroids to move for all cell lines considered. This observation is substantiated by the quantification of the protrusive over bulk projected areas plotted in Fig. 5d and with the measurements of movement, shown in Fig. 5e. Notably, no aggregation events are visible following actin inhibition. Taken together these observations confirm the prominent role of actin polymerisation and dynamics in the aggregation process.

While a substantial role for actin polymerisation was to be expected, given the phenomenological observations, we wondered whether we could see any effect relative to myosin dependent contractility, which has been shown to be a relevant feature in many collective migration models [37-39].

Our data indicate that the inhibition of myosin contractility and, consistently, of MLC activation through the inhibition of ROCK, dramatically impacts the compactness of the spheroids, allowing single cells or at most chains of cells to come out of spheroids, leaving the position of the spheroid bulk unaltered, as shown in Fig. 5c, bottom row. This behaviour is corroborated by the quantification of the protrusive over bulk projected area which is much higher than in the control, and with the quantification of movement, which is comparable to control if not higher (Fig. 5e). These results are consistent with previously reported observations on the role of myosin in collective migration and represent an interesting molecular insight on the ability of cancer cells to perform collective versus mesenchymal migration depending on the regulation of myosin contractility.

### CDA is perturbed by interfering with the signalling of PI3K/AKT/mTOR and MEK/ERK pathways

A fundamental question for the aggregation process and its driving molecular mechanism is whether this phenomenon is compatible with the signalling of upstream receptor-ligand dynamics or whether it is molecularly driven by purely cytoskeletal dynamics. It is indeed hard to find a unique molecular mediator common to all cell lines which evidently have profoundly different genetic alterations and behaviours. In order to address this question, we inhibited the most important signalling pathways downstream of cell surface receptor signalling, namely PI3K/AKT/mTOR and the MAPK/ERK pathways in order to check the impact of such perturbations with the aggregation phenomenon.

Due to the known involvement of such pathways in cell proliferation, in order to dissect the effect of inhibition on aggregation, we performed the experiments both starting from single cells and from preformed spheroids.

Single cell experiments (reported in Fig. 6a) showed that each inhibitor impacts both proliferation and aggregation, to different extents, as expected. The inhibition of MEK (AZD-6244, Selumetinib) has a clear effect on both aggregation and proliferation in MDA-MB-231 and MG-63 but not in PC-3, where it is essentially not effective.

**Figure 6:**
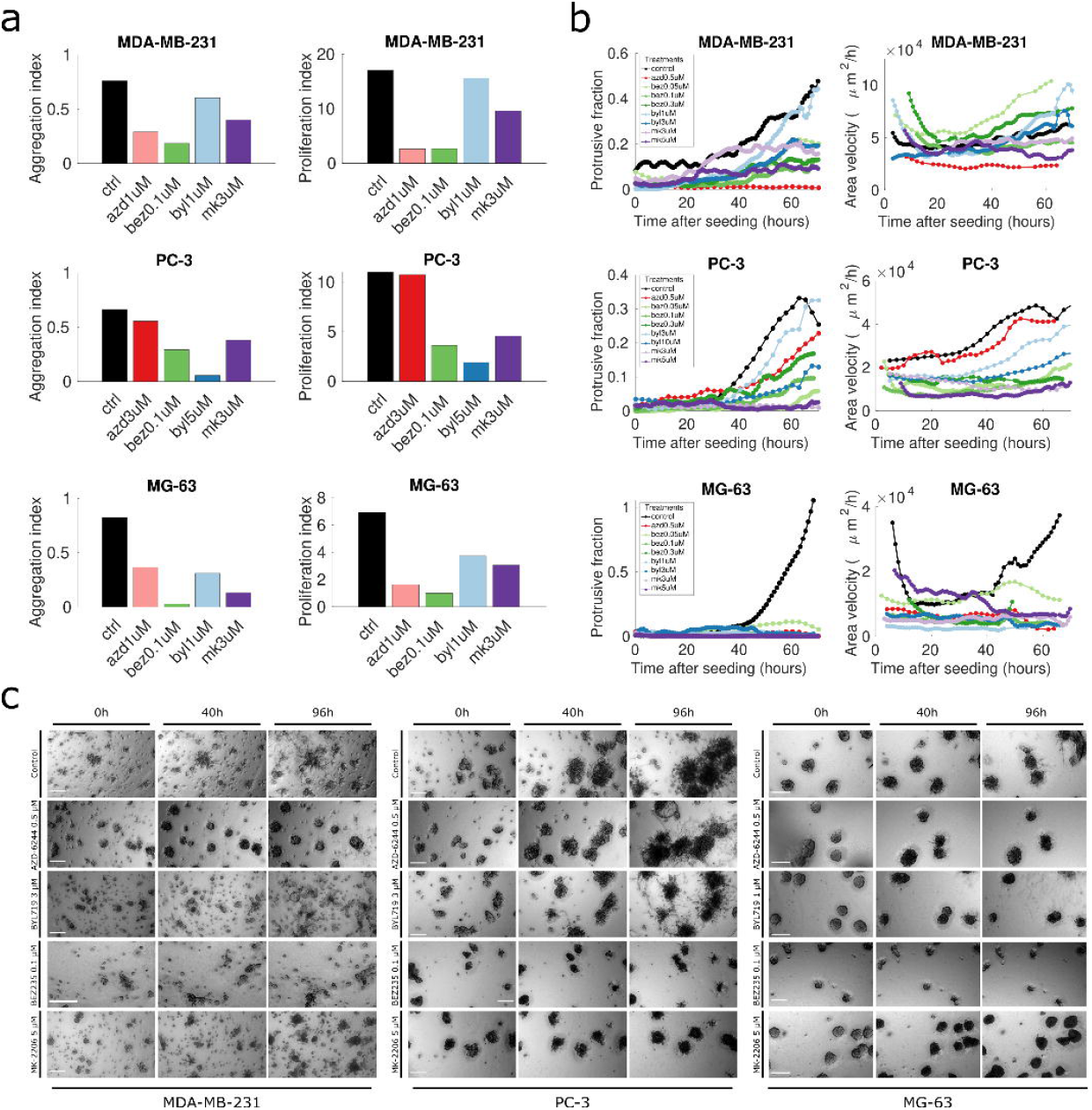
inhibition of downstream effectors MEK, AKT, PI3K and mTOR affects CDA to different extents. **(a1)** An aggregation index (equivalent to the density fraction shown in Fig. 2d and e) is shown for each inhibitor and each cell line to indicate the impact of each inhibitor on the aggregation process in each cell line. Aggregation assays are started from single cells. **(a2)** A proliferation index (obtained as the ratio between the total area of the spheroids in the images at 15 days and at 3 days after seeding) is shown for each inhibitor and each cell line in order to indicate the impact of each inhibitor on proliferation. Top to bottom: MDA-MB-231, PC-3 and MG-63. **(b1)** The ratio of protrusion to bulk regions in the images for each of the treatments and for the three cell lines is shown. **(b2)** Area velocity extracted from images as detailed in the Material and Methods section. Each colour represents a different treatment as detailed in the legend. **(c)** Pre-formed spheroids seeded in Matrigel were imaged for 4 days. Representative pictures at three time-points (0, 40 and 96 hours) for control and for each inhibitor are shown. Each of the three panels of images corresponds to a cell line; from left to right: MDA-MB-231, PC-3 and MG-63. Each row corresponds to one experimental condition, as reported on the left of each panel; from top to bottom: untreated, MEK inhibitor AZD-6244 0.5 μM, PI3K inhibitor BYL719 1 μM (MG-63) or 3 μM (MDA-MB-231, PC-3), PI3K-mTOR inhibitor BEZ235 0.1 μM, AKT inhibitor MK-2206 5 μM. Scale bar: 200 μm.

The PI3K/mTOR inhibition (BEZ235, Dactolisib) causes a reduction of proliferation and aggregation in all the three cell lines, while the pure PI3K inhibition (BYL719, Alpelisib) has only a modest effect on both proliferation and aggregation in MDA-MB-231, while it impacts more proliferation than aggregation in PC-3 and is effective on both aspects for MG-63.

Strikingly, we found the AKT inhibitor MK-2206 to have the most uniform effect on all three cell lines, acting both on aggregation and proliferation, especially at the highest concentration used.

In general, we found that none of the treatments had a clear effect on aggregation unless bound to an effect on cell proliferation, highlighting a dependence of one effect on the other. This is however consistent with the observation that aggregation is performed by multicellular aggregates, which when proliferation is blunt are much smaller. We found MG-63 to be sensitive to all treatments, while PC-3 tend to be less responsive to MEK inhibition and MDA-MB-231 less responsive to PI3K inhibition.

Results obtained by experiments made with pre-formed spheroids (Fig. 6b-c and SI Movie16-19) helped us to visualise a clearer effect on aggregation. To provide quantitative measurements of the behaviour of spheroids following the inhibitions, we employed the same indicators used in the previous section, i.e., the ratio between the protrusive and bulk projected areas and the area velocity. MEK inhibition is effective in impairing protrusion emission and movement in MDA-MB-231 and MG-63 but completely ineffective in PC-3. Conversely, PI3K inhibition had an intermediate effect on PC-3, a strong one in MG-63 and was ineffective on MDA-MB-231. Combined inhibition of PI3K and mTOR was effective in all three cell lines (slightly less on PC-3 as observed in single cell assays). AKT inhibition was the most effective in reducing migration and protrusion emission in all three cell lines. Quantitative results are consistent with what observed in time-lapse experiments shown in Fig. 6c, where effective treatments blunt aggregation, while intermediate effects correspond to partial aggregation of the spheroids. The results are therefore consistent with those obtained by single cell assays and show that both the protrusive fraction and the movement of spheroids are reduced or blunted by interfering with the signalling mediators MEK, AKT, PI3K/mTOR. Our results establish a common role for AKT in directional collective migration. The effect of the inhibition of PI3K and MAPK in PC-3 and MDA-MB-231 is instead cell line dependent, and consistent with the respective genetic backgrounds (PTEN deletion for PC-3 and KRAS and BRAF mutations for MDA-MB-231).

Taken together, these data support the hypothesis that aggregation is mediated by upstream signalling which then reflects into cytoskeletal dynamics, even though cell line specific molecular mechanisms need further investigation. This behaviour is particularly relevant given the conspicuous differences among the three analysed cell lines, i.e., their origin, the type of tumours, their phenotype and their genetic features.

### CDA is associated to the secretion of autocrine soluble cues in the medium

Previous results in our laboratory indicated that directional collective migration is consistent with the secretion of a soluble factor in the medium, which would stimulate the directional migration along concentration gradients, i.e., collective chemotaxis [32].

To further verify the consistency of the hypothesis of an autocrine loop mediating directional migration driving CDA, we tested the capability of all cell lines to migrate towards their own conditioned media. Conditioned media for each cell line were collected in serum-deprived 2D cultures at 24, 48 and 72 hours of culture and used in a classical chemotaxis assay (Transwell). Results show that conditioned media are all chemoattractant for cells, to different extents depending on the cell line (Fig. 7a,b). We found the chemotactic effect to be higher for later collection times for the majority of the cell lines, peaking at 48 or 72 hours of culture, suggesting that cell produced soluble cues are accumulated over time with degradation rates longer than 72 hours. Notably, MIA PaCa-2, picked among the non-aggregating cell lines, did not show significant attraction towards own conditioned media, except for a very slight effect of the conditioned media collected at 24 hours (SI Fig. 7a).

**Figure 7:**
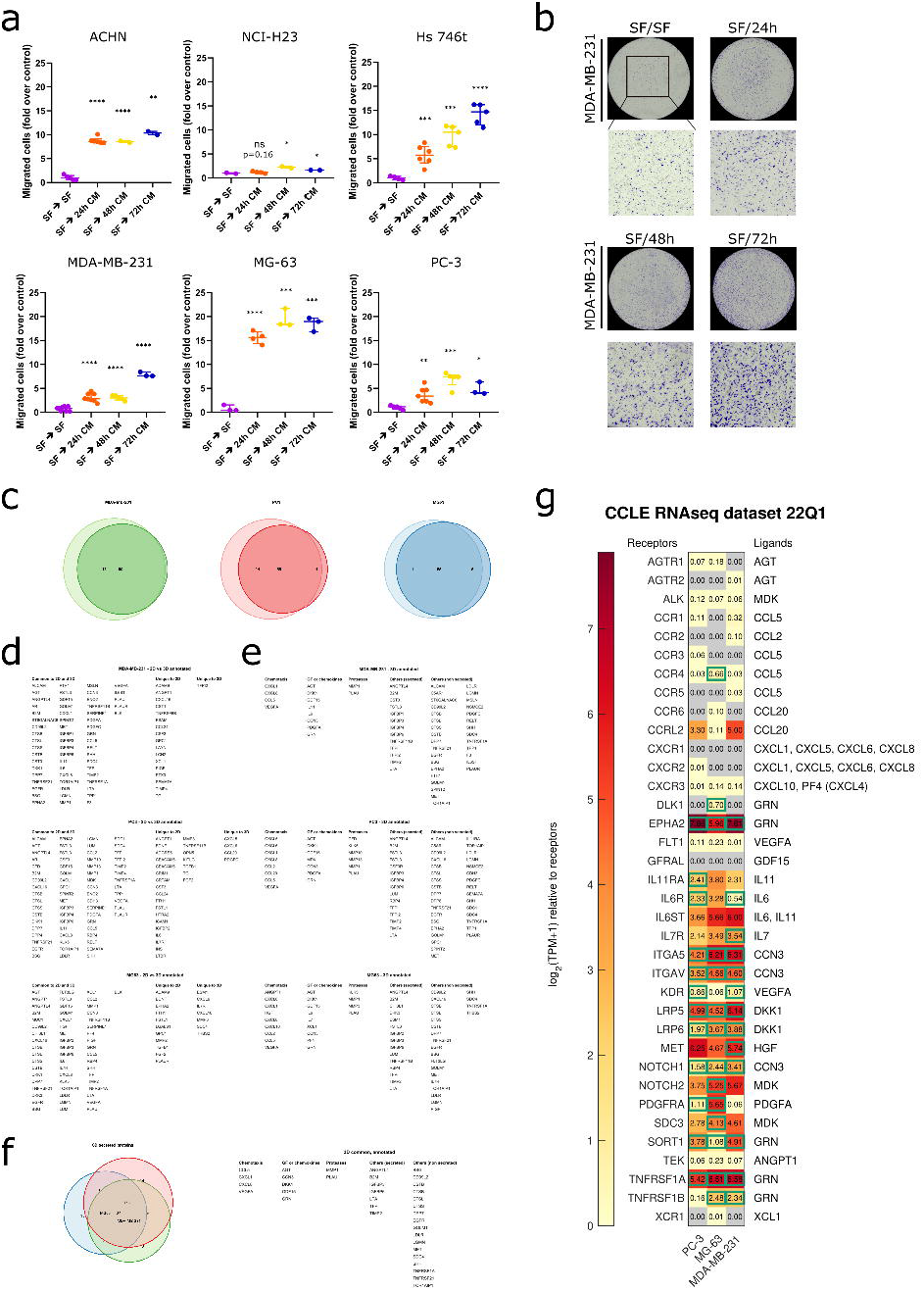
Conditioned media is chemoattractant for aggregating cell lines. **(a)** The plots report the migrated cells (fold over control) for different cell lines and conditioned media collected at different times. Each point on the plot represents data coming from a whole membrane. Migration of cells from serum free media towards serum free media (SF - SF) was used to normalise data as a control. Data are reported as median (horizontal line) with interquartile range. Statistical significance was assessed by performing a parametric one-tailed t-test with Welch’s correction (unpaired); *= P ≤ 0.05; **= P ≤ 0.01; ***= P ≤ 0.001; ****= P ≤ 0.0001. **(b)** Representative images of membranes corresponding to the experimental conditions reported in the graphs are shown. **(c)** Euler-Venn diagrams of factors that can be found in the corresponding conditioned media for 2D vs 3D, where proteins found in the controls with no cells, in the Matrigel or in the FBS were filtered out. **(d)** List of gene symbols corresponding to the proteins found in each part of the diagram. **(e)** List of gene symbols of proteins found in the 3D conditioned media, classified according to their gene ontology features. **(f)** Euler-Venn diagram of the three conditioned media and corresponding list with the common proteins annotated. **(g)** Heatmap of the transcriptional expression (CCLE) of the cognate receptors of all chemokines and growth factors for the three cell lines. Boxed values represent cases in which both the ligand and the receptor are expressed.

A purely chemokinetic effect of conditioned media was excluded by performing the same experiments with the conditioned media in the upper compartment (results are reported in SI Fig. 7b).

These data indicate that all the 6 cell lines secrete chemotactic factors in the culture medium.

In order to further deepen our analysis of the conditioned media content, we performed an antibody-based screening of 640 cytokines and growth factors for a series of experimental points. First, we ought to determine whether traditional 2D chemotaxis assays were comparable in terms of secreted proteins to the content of 3D conditioned media. To this aim we tested the content of conditioned media of three cell lines (MDA-MB-231, PC-3 and MG-63) in 2D and compared it with plain, serum deprived, cell-free media. Similarly, we screened the content of the conditioned medium of a 3D aggregation assay and compared it to cell-free serum containing media and cell-free matrix containing media, in order to select only the components due to autocrine secretion. The results of our analysis are shown in Fig. 7c-g. Our data indicate that the ensemble of secreted proteins in 2D and 3D is very similar, with more than 70% common secreted factors being present in both setups. All factors secreted by all the 3 cell lines and compared within 2D and 3D essays are listed in Fig. 7c.

We then characterised the secreted factors in the 3D assay. To this aim the list of all proteins was filtered eliminating the contribution of serum and cell-free Matrigel and then used to interrogate the database Uniprot [64] for gene ontology terms. We identified secreted proteins related to chemotaxis, growth factors or chemokines, proteases and other, both secreted and non-secreted, found proteins. The annotated list of these factors is shown in Fig. 7d. Our results indicate that all the three cell lines tested have both distinctive (i.e. line specific) and common factors, as shown in Fig 7e,f. Our data is therefore consistent with the hypothesis that CDA is dependent on secreted components.

We then screened the Cancer Cell Line Encyclopedia database [65] in order to establish whether receptors of the found cytokines and chemokines were expressed at least at the transcriptional level, and further supported these observations with a directed bibliographic search (see SI Table 2). We could identify 18 autocrine ligand-receptor couples that could be responsible for CDA. A few of these are shared among the three analysed cell lines, while others are cell line specific. Future studies will be required in order to rule out or confirm an involvement in CDA.

Taken together, our findings reinforce and confirm our interpretation of the data and are consistent with the hypothesis that aggregation is mediated by a gradient of diffusible factors.

### CDA mediates the formation of heteroclonal cell aggregates

We have shown that cells with a similar or identical genetic background, as those found in a single cell line, can be brought together by CDA. This aspect might potentially be of impact for phenotypic heterogeneity. An even more relevant question along this line is whether such a general mechanism might be able to bring together cells with a profoundly different genetic background, such as for example normal and tumour cells or even clonally distinct tumour cells.

To address this question, we performed aggregation assays with different cell lines by mixing cells expressing different fluorescent markers. Cell nuclei were labelled with fluorescent Histone2b (H2B) constructs and pre-formed spheroids with different cell lines were generated and imaged by means of time-lapse microscopy for several days. Screenshots of three representative couples of cell lines are shown in Fig. 8a (See also SI Movie20-22). Our results clearly show that spheroids of different cell lines form heterotypic clusters by performing directional migration. A representative example of the aggregation of heterotypic aggregates is reported in Fig. 8b-c. Multi-component, i.e., heteroclonal, spheroids are able to migrate collectively and to attract other (both homo-and hetero-typic) spheroids through the emission of multiple cells outgrowths similar to those found in homotypic aggregation events.

**Figure 8:**
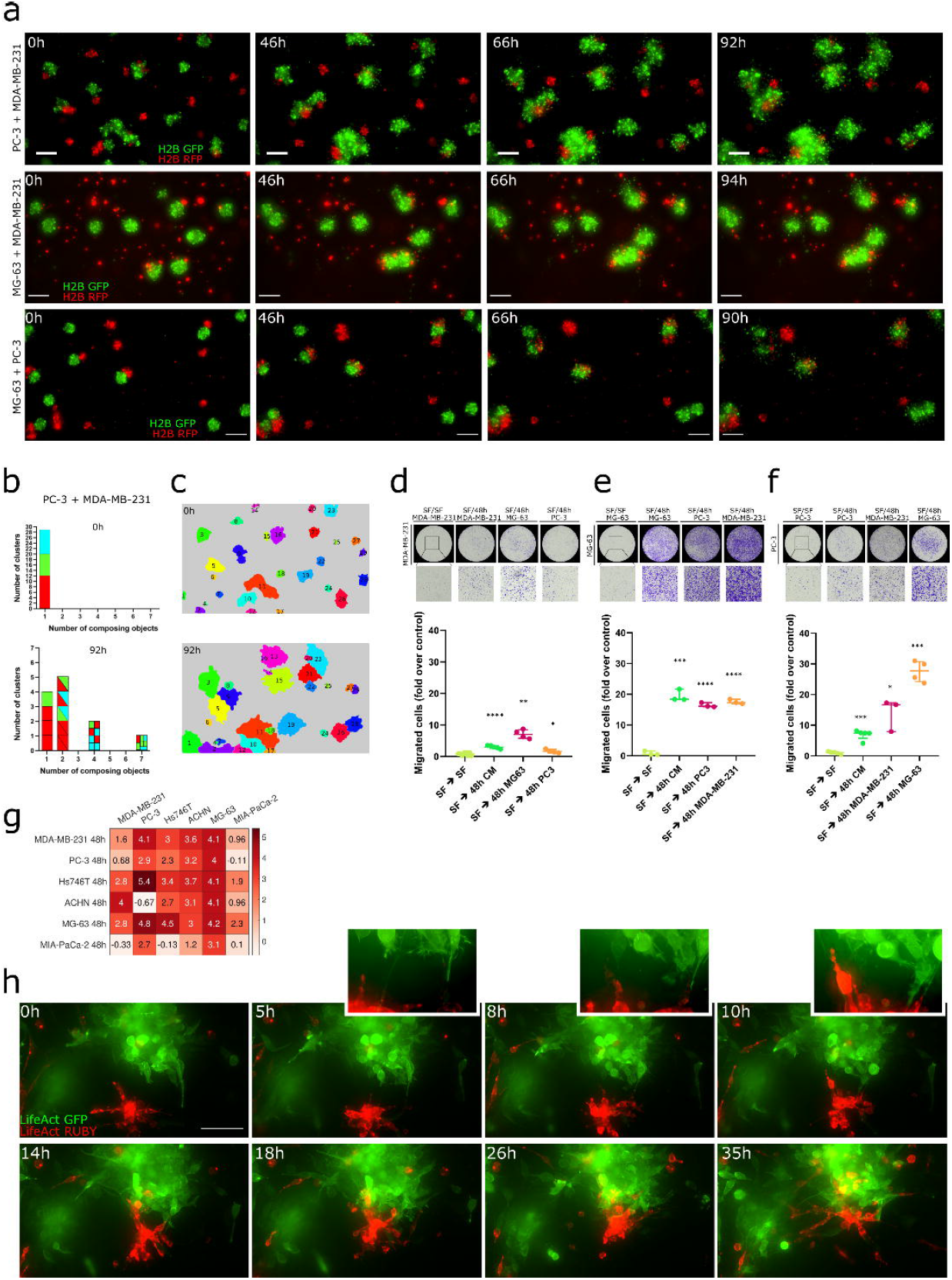
Heterotypic CDA is observed across different cell lines. **(a)** Snapshots of time-lapse experiments performed with pre-formed spheroids of cell lines couples expressing H2B-RFP (red) or H2B-GFP (green) were seeded in Matrigel. Each row represents (from the top to bottom): H2B-RFP MDA-MB-231 and H2B-GFP PC-3; H2B-RFP MDA-MB-231 and H2B-GFP MG-63; H2B-GFP MG-63 and H2B-RFP PC-3. Scale bar: 200 μm. **(b)** All the graphs and images in this panel refer to the aggregation assay between PC-3 and MDA-MB-231 (top-row images in a). The bar plots show the histogram of the number of composing aggregates of each object for the first (top, 0h) and the last (bottom, 92h) time-frame of the movie for the top row of panel (a), top row. Composing aggregates at frame 1 are conventionally considered as single objects as their aggregation happened before the assay started. **(c)** Images show the segmentation of the first and the last frames of the aggregation assay shown in panel (a), top row. Each object is identified by a numeric label and a unique colour in order to appreciate the aggregation of initially separated objects. **(d**,**e**,**f)** Chemotaxis assays (Transwell) performed with the cell line couples represented in panel (a). The plots report the migrated cells (fold over control) of different experiments. Each point on the plot represents a whole membrane. Data are reported as median (horizontal line) with interquartile range. Statistical significance was assessed by performing a parametric one-tailed t-test with Welch’s correction (unpaired); *= P ≤ 0.05; **= P ≤ 0.01; ***= P ≤ 0.001; ****= P ≤ 0.0001. Representative membranes corresponding to the experimental condition reported in the plots are shown on top of each plot. **(g)** The matrix reports the migrated cells (fold over control) of different Transwell assay experiments performed to test whether the conditioned medium of each of the 6 aggregating cell lines act as chemoattractant for all the other cell lines. Indicated values correspond to the log2 of the median fold-change. Experiments were conducted by seeding onto the Transwell membrane cell lines listed in the column labels and by adding in the lower compartment 48h conditioned media collected from cell lines listed in the row labels. **(h)** Representative images of time-lapse movies showing an aggregation event between pre-formed spheroids of LifeAct-Ruby (red) MDA-MB-231 and LifeAct-GFP (green) PC-3 cells in Matrigel. The section of the images corresponding to the area between the two clusters was enlarged to facilitate the visualisation of the protrusions and it is shown in the top-right inserts. Time labels indicate hours after seeding. Scale bar: 100 μm.

A necessary underlying condition to the observed behaviour is that conditioned media are able to act as chemoattractant also for other cell lines, notably for those that are involved in heteroclonal aggregation. To verify this hypothesis, we performed Transwell assays with aggregating cell lines and used conditioned media originating from different cell lines, as shown in Fig. 8d-g (See also SI Fig. 8a). Our results indicate that the conditioned medium of each of the 6 cell lines act as chemoattractant, to different extents, for all the other cell lines. In particular, the conditioned media collected from the cell line MG-63, which is the cell line with the highest percentage of aggregating cells (as reported in Fig. 1f), is chemoattractant for all the remaining 5 cell lines and has also the greater chemotactic capability, in terms of the number of migrated cells over control, in comparison with conditioned media of other cell lines. By contrast, the conditioned media of MIA PaCa-2, picked among the non-aggregating cell lines, is much less effective (green experimental points in the graphs of SI 8a). Interestingly, the conditioned media collected from MDA-MB-231, MG-63 and ACHN is chemoattractant for MIA PaCa-2 (Fig. 8g and SI Fig. 8b). In order to understand whether subcellular protrusions can be found in heterotypic aggregation as well, we performed time-lapse experiments at high spatio-temporal resolution with pre-formed spheroids made with two different cell lines expressing fluorescent LifeAct. As in the case of homotypic aggregation, we observed that the coalescence between two distinct, initially separate, spheroids of PC-3 and MDA-MB-231 is accompanied by the emission of protrusions, as shown in Fig. 8h (and Movie23). These data confirm the hypothesis that CDA can drive the formation of heteroclonal aggregates.

## DISCUSSION

Our study provides a novel 3D model to investigate cancer cell collective migration dynamics and identifies such behaviour as the mechanism driving homo- and hetero-typic aggregation of cancer cell clusters. We demonstrate that the movement of cells and clusters is directional and mediated by single and multicellular actin-rich protrusions and that molecular inhibition of cytoskeletal dynamics results in impaired CDA. Our results pinpoint a crucial role of myosin II in the retainment of cell-cell contact during cancer collective migration as observed in other biological contexts [37-39]. Our data support the notion that, while CDA does not require cell proliferation directly to occur, the two phenomena are coupled, i.e., faster proliferation enhances CDA. We hypothesise that cluster-cluster interaction at distance is mediated by soluble cues, such as for example the autocrine secretion of chemoattractants, which would in turn generate spatial gradients mediating directional migration. This hypothesis is supported by the capability of cells to migrate toward their own conditioned media and by the finding that conditioned media collected from CDA assays contain a significant number of secreted proteins, many of which involved in cytokine, growth factor or even chemotactic activity. Notably, a subset of such ligands is associated to transcriptionally active cognate receptors. Consistently, CDA is impaired when cell surface receptors downstream effectors are inhibited. This result also indicates that aggregation is not purely driven by cytoskeletal dynamics but is indeed compatible with upstream receptor-ligand signalling. Interestingly, we observed reciprocal aggregation between cells originally derived from different tissues, identifying CDA as a potential mechanism at the basis of the formation of heterogeneous tumours.

The role of chemotaxis in cancer dissemination is documented [66], but mostly involving paracrine mechanisms, e.g., between tumour cells and normal cells [19-23, 67]. Nevertheless, autocrine loops are widespread across cancer cells [68-74] and chemokines and chemokine receptors are often overexpressed in both metastatic and primary tumour cells, but not in the corresponding normal counterpart [75]. We speculate that expression of such receptors or ligands might determine a higher fitness of cancer cells in comparison to other cellular populations within the primary tumour site, thanks to ligands acting both as chemotactic factors and as mitogens. This would also imply a role for CDA as a self-sustaining mechanism by which cancer cells could augment their survival rate.

It has indeed been proposed that greater metastatic capability of CTCs is due to proliferative signals that cells send to each other to improve their survival rates [17]. In light of our findings, we can suppose that such proliferative signals might coincide with (or be associated to) chemotactic signals, conferring a double advantage to CTCs. Remarkably, there is one out of 6 cell lines which we reported to perform CDA deriving from a primary tumour site, suggesting that the capability to migrate in response to chemical cues might in principle be relevant in the primary tumour site, in the dissemination and possibly in the metastasis formation. On the other hand, we anecdotally report that the frequency of cell lines derived from metastatic sites is lower in the non-aggregating category, hinting at the conclusion that CDA might be more diffused in metastatic cancer cells.

The observation that CDA still proceeds, even in proliferation defective cells, has an important therapeutically relevant consequence. Tumour cells treated with drugs targeting cycling cells or proliferation directly will not stop performing CDA. Therefore, cytostatic drugs would not affect the ability of tumours to evolve towards metastasisation, or to increase local tumour cell density to increase survival probability and metastasisation.

The observation that cells originating from spatially distant heterogeneous clones can merge by collective directional migration might be relevant in the context of tumour dissemination as well, as cells expressing even only a receptor for a ligand autocrinely secreted by a separate group of tumour cells, might be attracted during their route from the primary site to distant organs, as already reported for interaction between tumour and normal cells. Such a mechanism might provide a possible explanation of the early acquisition of clonal heterogeneity within the tumour, which represents a big obstacle in the development of effective pharmacological treatments [76].

Our findings are among the few to describe directional migration in collective migration [30, 32, 34, 77]. As already pointed out by previous works, collective chemotaxis might have important differences to single cell chemotaxis in terms of sensing. The detailed mechanisms of transmission of the information on orientation across cells in a multicellular aggregate might rely on purely mechanical signalling among the cells or on chemical intercellular signalling. While these aspects are almost entirely to be elucidated, the dependence of CDA on concentrations and on concentration gradients might be completely different than for single cells, and might result in different dynamical behaviours.

While it has to be expected that the molecular mediators of such phenomenon are not unique, we acknowledge that more detailed molecular studies are required to further investigate the role of such a phenomenon *in vivo* as well as to better understand the role of other concurrent mechanisms such as mechanical forces and specific proteolytic activity of tumour cells. Further investigations might help elucidate if and when CDA is relevant during cancer progression and whether there are any significant correlated effects on prognosis.

Notably, the results obtained by inhibiting the signalling pathway effectors might indicate somewhat common mechanisms across cell lines and therefore among different tumours, which, considering the extreme diversity between the analysed cell lines, is an interesting unexpected finding and point at CDA as a rather universal mechanism rather than as a specific aspect of a single cell line or a single molecular mediator.

Our study describes CDA as a novel widespread phenotype and paves the way to further investigation about the biology of such phenomenon and the potential advantage that CDA might have in tumour progression.

## ACKNOWLEDGMENTS

This study was supported by funding from “Associazione Italiana Ricerca sul Cancro” MFAG-2020 n. 25040 to AP and IG-23211 to LP; University of Torino Fondo Ricerca Locale 2019 PULA_RILO_19_01 to AP; FPO-Young Investigator Grant 2020 to AP; Fondazione Umberto Veronesi postdoctoral fellowship 2017 and 2018 to AP. We thank Dr. E. Vallariello and Prof. F. Bussolino for kindly providing CFPAC-1, PANC-1 and MIA PaCa-2 cell lines, Ms. I. Cantarosso for helpful assistance with *in vitro* experiments, Dr. R. Albano for the Cell Culture Facility service and Dr. L. Fontani for the Gene Transfer Technology Facility service.

## MATERIALS AND METHODS

### Cell culture

The cancer cell lines used are listed in SI Table1 and were purchased from ATCC. Culture media (all purchased from Sigma-Aldrich), unless specified, were supplemented with 10% Fetal Bovine Serum (ThermoFisher Scientific), 200 U/ml of penicillin, 200 μg/ml streptomycin (Sigma-Aldrich) and 2Mm L-Glutamine (Sigma-Aldrich). Cells were kept at 37 °C under 5% CO2 humidified atmosphere. Cells were counted automatically by using TC20 Automated Cell Counter (BioRad). Medium was replaced three times a week. Each cell line was used up to passage 15. PCR-based Mycoplasma test was performed once a week.

Aggregation assays either starting from single cells or from spheroids were carried out in standard tissue culture plastic dishes (Falcon), multi-well plates (Corning) or imaging dedicated supports (glass-bottom 35 mm dish, Ibidi 81218-200; 96 well-plates Corning; μ-Slide 8 Well Chamber Slide, Ibidi 80827; 2-well silicone inserts, Ibidi 81176).

### Cancer cell spheroid formation

Spheroids from cancer cell lines were generated by using the hanging drop technique. The protocol was adapted from the procedures reported in [78, 79]. A methylcellulose stock solution was prepared by dissolving 6 grams of autoclaved methylcellulose powder (M0512 Sigma-Aldrich) into 250 ml of preheated (60°C) serum free culture medium, with the help of a sterile magnetic stirrer, for 2h. 250 ml of 20% FBS culture medium with penicillin/streptomycin was added to a final volume of 500 ml, to obtain a final concentration of 10% FBS, and the solution was mixed overnight at 4°C. The 500 ml were aliquoted and centrifuged at 5000g for 2h at room temperature. The supernatant was collected and used for the spheroid assay, while the pellet was discarded. For spheroid generation, cells were detached, counted and mixed with 80% culture medium and 20% methylcellulose stock solution to obtain a final concentration of 0.24% methylcellulose. Drops of 30 μl were generated by using a multichannel pipette and laid down onto the inner part of a 150-mm petri dish lid. The lid was carefully reversed to let the cells growth in the hanging drops, under non-adherent culture conditions. Cells were maintained humidified by adding PBS into the plates and kept in the incubator. After three days, spheroids were carefully harvested by washing out the drops with PBS and collecting the solution into a 50 ml tubes. Spheroids were gently spin down at 500 RPM for 5 minutes, the supernatant was removed and spheroids were mixed with Growth Factor Reduced (GFR) Matrigel (Corning) by using wide-bore (and precooled) tips to avoid desegregation. Spheroids pre-mixed with Matrigel were seeded as described in the following paragraph.

The following number of cells per drop was used to form spheroids of about 200 μm diameter: ACHN: 100; CFPAC-1: 300; Hs 746T: 100; NCI-H23: 100; MG-63: 100; MDA-MB-231: 300; PC-3: 50; MIA PaCa-2: 300.

### 3D cell and spheroid culture

-Matrigel and BME: For 3D cell culture, cells and pre-aggregated spheroids were mixed at the desired density with GFR-Matrigel (Matrigel® Matrix, Corning) diluted with culture media to a final concentration of 8 mg/ml. To avoid the bi-dimensional growth of cells or spheroids underneath (on the plastic bottom of the plates) or on top of the GFR-Matrigel during the 3 week-long assays, a “sandwich” of three layers was prepared as follow: a first layer of GFR-Matrigel was deposed on the bottom of 96- or 48-well plates. A second layer of GFR-Matrigel plus cells or spheroids was added on the top of the previous layer and a third layer of GFR-Matrigel was deposed on top. Each layer was left to polymerise for 10 minutes at 37°C before adding the next layer. Plates were kept on ice to depose the first layer and GFR-Matrigel was manipulated in ice by using precooled tips, to avoid hydrogel solidification. After polymerisation, wells were filled with culture media (200 μl in 96 well-plates and 500 μl in 48 well-plates) and plates were kept at 37°C. Culture media were replaced three times a week. The same procedure was followed to perform aggregation assays whit Cultrex BME (CultrexTM Basement Membrane Extract, R&D).

The following volumes of Matrigel were used for each of the three layers depending on the culture supports: 96-well plates: 30 μml– 40 μl (including cells) – 30 μl; 48-well plates: 80 μl – 100 μl (including cells) – 80 μl; glass bottom 8 slides: only one layer of 80 μl, to obtain a sufficiently-thin slice suitable for microscopy experiment with short working distance objectives (suitable only for one or two day-long experiments); 2-well silicone inserts: only one layer of 40 μl (same reason).

-Rat Collagen: Type I rat tail Collagen (Roche, Manheim, Germany) was dissolved in 0.2% acetic acid to a final concentration of 3 mg/ml. To induce polymerisation, Collagen was mixed with chilled 1 M NaOH and 10X culture medium according to the ratio: 1:0.032:0.1 (vol/vol). The pH of the collagen solution was adjusted with NaOH to reach pH between 7.0 and 7.5 (salmon pink colour by eye). The neutralised collagen solution was then incubated on ice for 1 h to increase the viscosity. A first layer of collagen solution was plated o the bottom of the wells. After polymerisation (45 minutes at 37°C), a second layer of Collagen mixed with cells was added on the top of the previous layer and let gelling 45 minutes at 37°C. Wells were then filled with culture media.

-PuraMatrix: before starting the experiments, the stock solution (PuraMatrix™ Peptide Hydrogel, 1% w/v, Corning) was shaken at room temperature to decrease the viscosity. To embed spheroids in PuraMatrix, after centrifugation, spheroids were washed twice with a 10% sucrose solution and the pellet was carefully resuspended in PuraMatrix mixed with the same volume of 20% sucrose solution. The spheroid/hydrogel mixture was carefully dispensed along the side of the well. In order to promote gelation, culture medium was very gently added on the top of the hydrogel.

Medium was replaced every 30 minutes for three times to allow complete gelation.

### Drug treatments

To test the effect of pharmacological inhibition, cells and spheroids were seeded in Matrigel as described above and media were replaced with media containing the inhibitors at the desired concentration diluted in DMSO. In the case of single cells, inhibitors were added two days after seeding, while in the case of pre-aggregated spheroids, inhibitors were added a few hours after seeding. Media were replaced three times a week. For actin perturbation we used (Fig. 4) 1 μM Latrunculin A (Sigma-Aldrich); 1 μM Cytochalasin D (Sigma-Aldrich); 100 μM CK666 (Sigma-Aldrich);10 μM Wiskostatin (Sigma-Aldrich). For Myosin perturbation we used: 100 μM Blebbistatin (Calbiochem); 20 μM Y-27632 (Sigma-Aldrich).

For the results shown in Fig. 5, the following inhibitors were used PI3K/mTOR inhibitor BEZ235 (Selleckchem) 50 nM, 100 nM and 300 nM; PI3K inhibitor BYL719 (MedChemExpress) 1 μM, 3 μM, 5 μM and 10 μM; AKT inhibitor MK-2206 dihydrochloride (MedChemExpress) 1 μM, 3 μM and 5 μM; MEK inhibitor AZD-6244 (MedChemExpress) 0,5 μM, 1 μM and 3 Mm.

To test the effect of mitomycin cells were plated in 96 wells Ibidi Imaging plates one day before starting the experiment at the density of 4000 cells/well. In the following days the cells were treated with mitomycin at 0.3μg/ml, 0.5 μg/ml, 0.75 μg/ml, 1 μg/ml, 1.5 μg/ml and 2 μg/ml for 3h, 6h or 24h. Cell cycle blocking was assessed using the Click-iT EdU Imaging Kit (Invitrogen). Cells were washed twice with PBS and incubated with EdU for 8 hours, 4% PFA-fixed, permeabilised with 0,5% Triton X-100, washed with 3% BSA and incubated with Click-iT reaction cocktail for 30 minutes at room temperature. Cells were then washed in 3% BSA and stained with DAPI.

### Chemotaxis assay

Chemotaxis was assessed by using 6.4 mm Transwell Permeable Support (Falcon) with 8-μm pore PET membrane inserts. 3 × 10^4^ cells (or 1 × 10^4^ in the case of MG-63 cell line) were seeded on the permeable membrane with 100 μl of medium. Cells were kept in the incubator over-night to allow the attachment. The following day, cells were gently washed three times with PBS and 400 μl of serum free medium was added. The low compartment was filled with 750 μl of conditioned media. After 24 hours of incubation, non-migrated cells on the upper surface of the membrane were removed with a cotton swab, while cells passed through the membrane were fixed with 2,5 % glutaraldehyde in PBS, washed 2x with PBS and stained with 0.1% crystal violet solution in methanol. Once dry, membranes were imaged on an inverted microscope and images were analysed by means of automatic segmentation as detailed in the Image Analysis section.

To obtain the conditioned media, 2 × 10^6^ cells were seeded into a 10-mm Petri dish with complete media and let attach overnight in the incubator. The following day, cells were washed with PBS and 2 ml of serum free media were added. Conditioned media were collected after 24, 48 and 72 hours and filtered with 0.22-μm filters before being used for the assay.

### Cytokine screening

Cytokine screening was performed with a glass slide-based antibody array (RayBiotech, GSH-CAA-640) containing 640 cytokines, growth factors and other secreted proteins. Sample preparation was performed according to the manufacturer’s instructions.

Samples were prepared as follow: conditioned media in 2D culture plates at 72-hour time-point were obtained as reported in the “Chemotaxis assay” section. Once collected, supernatants were immediately centrifuged at 2500 g for 10 minutes at 4°C and then stored at −80°C until use. Transwell assays were performed with the collected media in order to confirm activity. Supernatants from 3D aggregation assays were collected after 10 days of culture. Culture media were replaced three times a week and the last replacement was made 72 hours before harvesting. Media were collected either from the top of the gel or by mechanically disrupting the gel in order to check the secreted fraction trapped within the gel. Supernatants were centrifuged at 2500 g for 10 minutes at 4°C immediately after collection and then stored at −80°C until use.

For each condition we performed the matched controls (no cells for 2D conditioned media, medium with FBS, Matrigel with no cells, with or without mechanical disruption). Data were analysed in house by means of custom written Matlab codes. Briefly, a common threshold value was set by minimising the number of hits in cell free, serum free and Matrigel free media. Components exceeding this threshold coming from cell free controls (with or without serum and Matrigel) were filtered out. Protein content of serum and Matrigel themselves was reduced to minimum due to the use of human antibodies spotted on the slides. The list of proteins was first converted into gene symbols and relative gene ontology terms were fetched by interrogating Uniprot [64] by means of custom-made Python scripts. The list of candidate cognate receptors corresponding to secreted ligands was manually compiled by fetching the literature, and transcriptional presence (in the form of log2 of TPM) was fetched for the three cell lines from the Cancer Cell Line Encyclopedia [65].

### Plasmids and lentivirus production

Plasmids were purchased from Addgene. LV-GFP and LV-RFP were a gift from Elaine Fuchs (Addgene plasmid #25999 and Addgene plasmid #26001) (Beronja et al. 2010). pLenti.PGK.LifeAct-Ruby.W and pLenti.PGK.LifeAct-GFP.W were a gift from Rusty Lansford (Addgene plasmid #51009 and Addgene plasmid #51010). Lentiviruses were produced by calcium phosphate transfection of lentiviral plasmids together with packaging (pCMVdR8.74) and envelope (pMD2.G-VSVG) plasmids in 293T cells. Supernatant was harvested 24 and 48 h after transfection, filtered with 0.45-μm filters, precipitated (19,000 *g* for 2 h at 20°C), and suspended in PBS at a higher concentration. The multiplicity of infection (MOI) was determined by infecting HeLa cells and by quantifying GFP, RFP or Ruby-positive cells by flow cytometry. All cell lines were infected considering a MOI of 2 viral particles per cell, except MG-63 which required a MOI of 25 viral particles per cell. Positivity to fluorescent proteins was assessed 48 hours after infection.

### Imaging methods

Time-lapse experiments were performed on inverted microscopes, either confocal or widefield, equipped with a motorised stage and an incubator to keep the plate stably at 37° and 5% CO_2_. For 2 or 3 week-long time-lapse experiments, cells were kept in the incubator and imaged once a day. To observe the protrusion composition (fig 1c), infected cells (with fluorescent H2B and LifeAct) were used to form spheroids, as described above. Spheroids were embedded in Matrigel and after 2 to 3 days culture medium was removed, spheroids were washed twice with PBS and fixed with 4% PFA for 30 minutes. PFA was removed by washing three times with PBS, and samples were kept in PBS for subsequent imaging.

Images throughout the paper are obtained by a number of different techniques and instruments. Temporal grayscale series were obtained by the BioTek Cytation 3 equipped with a 4X objective (Olympus). Fluorescence images were obtained with either a confocal Leica SP8 equipped with dry 20X or immersion 40X or 60X or with a widefield microscope Nikon Ti2 (Lipsi) equipped with dry 20X 0.75, immersion 40X 1.15 or 60X 1.4 and a wide field of view monochromatic camera (IRIS 15, Photometrics). Live imaging was performed with an automatic water dispenser for immersion objectives. Fluorescence images were obtained by acquiring z-stacks which were then maximum intensity projected. Widefield fluorescence images were previously deconvoluted with Richardson-Lucy algorithm. In selected images, a Gamma correction was used to increase the contrast of the lower intensity protrusion with the respect to the body of spheroids for visualisation reasons. Original images are available upon request. Whole Transwell membranes were acquired by the same widefield microscope with an RGB camera by stitching several images together.

Grayscale images in timeseries are obtained by projecting stacks (typically 1mm thick) of single brightfield images taken every 50 μm and then projected onto a single image with extended depth of field (EDF) [80] based on the maximisation of the local image variance (with a kernel size of 8 μ m), implemented in Fiji through the plugin developed by EPFL [81]. Images were handled with custom-written scripts for Fiji or Matlab. Deconvolution for widefield images was performed in the case of EdU staining by means of NIS-Element software (Nikon Instruments) with traditional Richardson-Lucy algorithms and pre-measured PSF.

For assessing the effect of mitomycin cells were labeled with a live nuclear marker (NucBlue™ Live Cell Stain Thermo Fisher Scientific, Waltham, MA, USA) and imaged in the UV wavelength. To count the apoptotic events cells were treated with a fluorescent dye detecting Casp3/7 activation (CellEvent™ Caspase-3/7 Green Detection Reagent, Thermo Fisher Scientific, Waltham, MA, USA).

### Image analysis

In order to quantify the protrusive fraction of a given time-lapse series we implemented the following image analysis algorithm. A large set of timeseries was fed to a machine learning segmentation software (Ilastik [82, 83]) with different classes for background, bulk and protrusive areas. Multinary images obtained were then processed by a custom written Matlab algorithm to filter out small isolated structures (<50 μm radius) and calculate the ratio between the fraction of the protrusive area in the image to the total cluster occupied region at the first frame, in order to account for the variable number of spheroids in the field of view.

The fraction of aggregating cells was calculated by manually screening all cells appearing in the first frame of single-cell aggregation assays for aggregation events found anytime during the experiment. All cells at seeding were therefore subdivided into aggregating or not, forming a fraction.

The effect of drug treatments on single cells aggregation assays was quantified as indicated in the figure 5 caption.

The number or independent objects in EDF time series were counted and overlapped objects were checked in the original non projected stack. In order to obtain quantitative parameters, the number of objects over time was fitted to a sigmoid, shown in Fig. 2c.

The estimation of the velocity from EDF time series was performed as follows. First images were aligned by means of cross-correlation methods in order to correct for stage offset and gel large scale movement. PIV-based image alignment was implemented in Matlab, and details of the method can be found elsewhere [84]. After image alignment, segmented images were used to compute area differences at 3 hours in order to consider only relatively fast movements and not overall growth. Such differences between frames were used as an estimation of the velocity by dividing the corresponding area in squared microns by the time interval.

Transwell images were segmented by means of the previously mentioned software Ilastik and binary images were quantified by Matlab custom written scripts. EdU stained z-stacks were segmented with Ilastik, and positive cells were selected by setting a threshold computed from control images. Contrast in the panels shown in the manuscript in Fig. 4d has been set in the same fashion for all images. Apoptotic events counted in mitomycin-treated images were identified by calculating the number of events overtime (as green flashes) and summing up all the events to obtain a toxicity over the indicated period of time.

### Statistical and data analysis

The kinetic data of aggregation (Fig. 2a,b) were fitted to a sigmoid shown in Fig. 2c. To estimate the doubling times, we fitted the data extracted from the segmentation of single cells or spheroids, and corrected for the dependence on the radius. The total volume of the objects is proportional to the number of cells. For an initially monodisperse distribution the area elevated to the power 3/2 is a pseudo-volume, which can be used as a proxy for the total number of cells.

Therefore, to extract the approximated doubling time of cells we fitted the area with an exponential A*2^(t/tau) and calculated the doubling time as 2/3tau.

Statistical analysis on the data of chemotaxis assays was performed using GraphPad Prism 8.0.0 software. Statistical significance was assessed by performing a parametric one-tailed t-test with Welch’s correction (unpaired). P ≤ 0.05 was considered significant. When not significant, the p value has been indicated in the figure.

## Notes

https://osf.io/meygs/

### Competing Interest Statement

The authors have declared no competing interest.

