## Supplementary Material for "Collective directional migration drives the formation of heteroclonal cancer cell clusters"

### Supplementary Material of the paper: Collective directional migration drives the formation of heteroclonal cancer cell clusters

**Miriam Palmiero[1,2], Laura Di Blasio[1,2], Valentina Monica[1,2], Barbara Peracino[3], Luca Primo[1,2,*] and Alberto Puliafito[1,2,*]**

[1] Candiolo Cancer Institute, FPO - IRCCS, Str. Prov. 142, km 3.95, 10060 Candiolo, Italy.
[2] Department of Oncology, University of Turin, 10060 Candiolo, Italy
[3] Department of Clinical and Biological Sciences, University of Turin, San Luigi Hospital, 10043 Orbassano, Italy
[*] these authors contributed equally

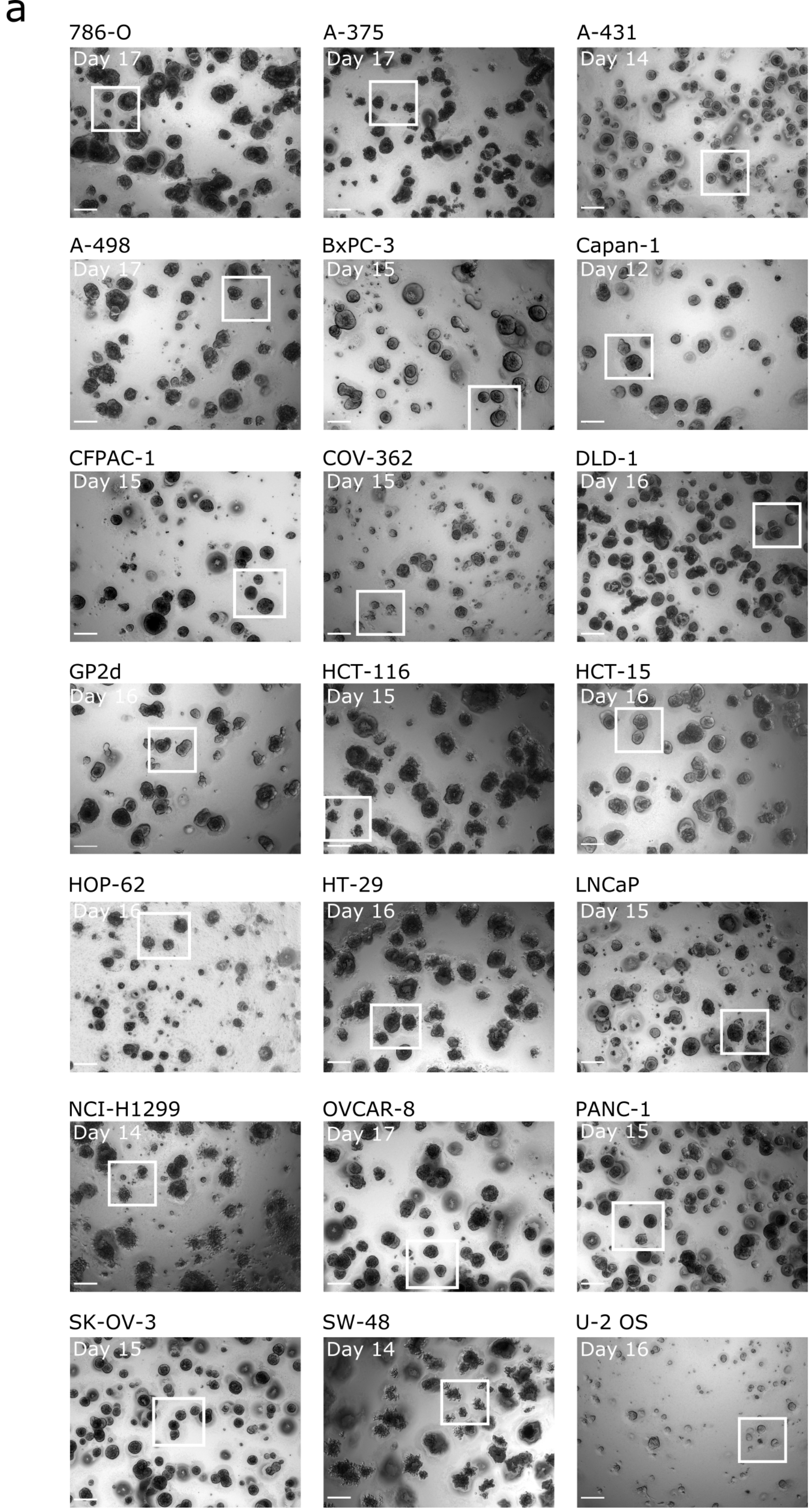


**Supplementary figure 1: (a)** Representative CC lines, derived from different tissues of origin, were used in our aggregation assay showing a non-aggregating phenotype. Cells were seeded as single-cell suspension in Matrigel and imaged by means of time-lapse bright-field microscopy for several weeks. One representative snapshot for each cell line at the end of the aggregation assays (time-points are indicated on the top-left corner of the pictures) is shown. White squares point out clusters that just grow without touching each other. Scale bar: 200 µm.

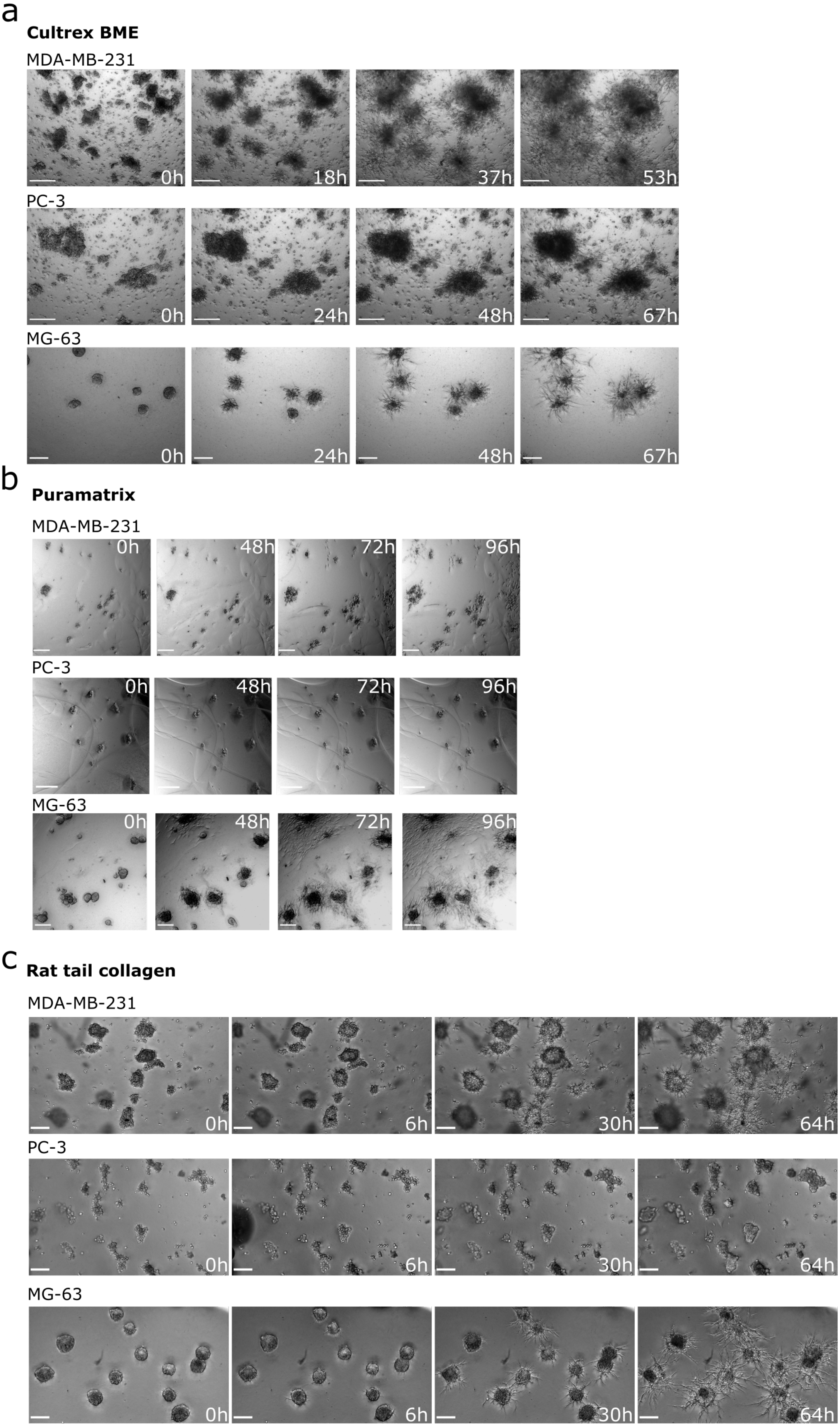


**Supplementary figure 1B:** Representative snapshots of preformed spheroids embedded in different hydrogels. **(a)** Cultrex BME. In this hydrogel pre-formed spheroids behave analogously to what observed in Matrigel. **(b)** Corning Puramatrix. Here we found spheroids to be much less protrusive and non-migratory. **(c)** Roche Type I Rat-tail Collagen. Spheroids grown in collagen are generally more protrusive, displaying multicellular outgrowths collectively invading the surrounding matrix. No bulk spheroid movement is observed in this case. Scale bar: 200 μm.

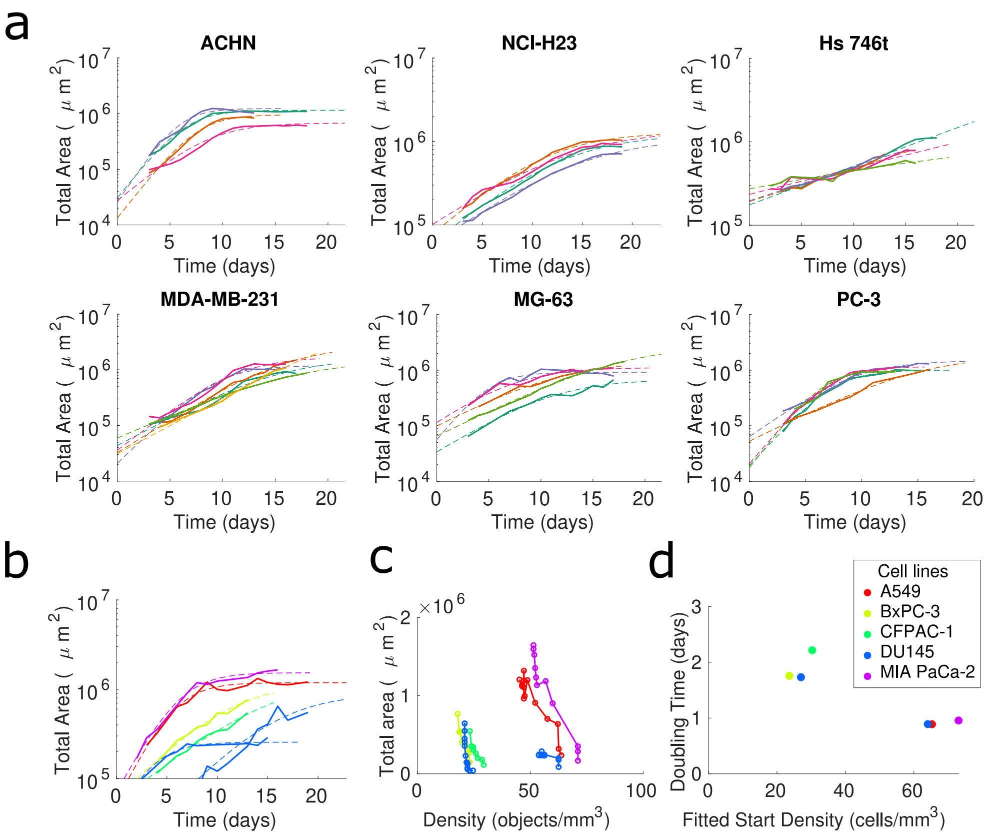


**Supplementary figure 2:(a,b)** Timeseries of the total area for each density of the indicated aggregating cell lines (a) and non-aggregating cell lines (b), indicated in the title of the plot or in the legend in panel (d). Dashed lines represent the fit with a saturating exponential. **(c)** Scatter-plot of the total area vs density for non-aggregating cell lines. **(d)** Doubling times for non-aggregating cell lines plotted against the starting density.

a

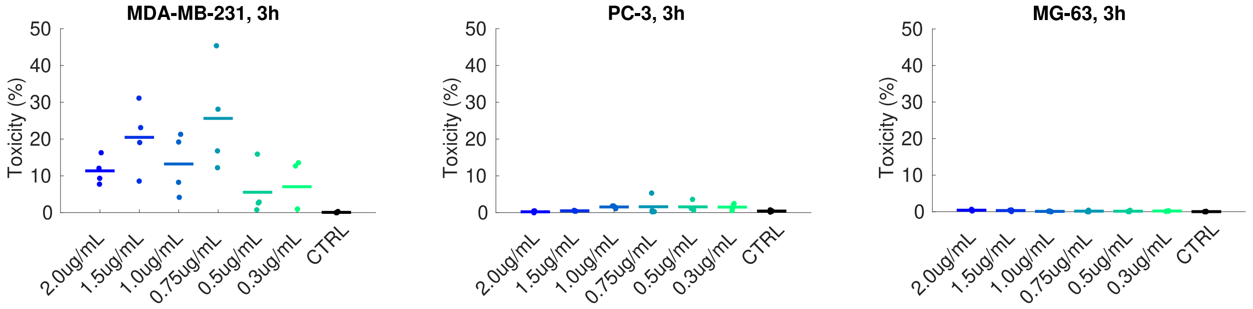


**Supplementary figure 4: (a)** The level of toxicity for each concentration of mitomycin is evaluated by assessing the cumulative number of apoptotic events normalized by the number of nuclei at the beginning of the experiment, marked by a fluorescent signal triggered by Cell Event Casp3/7 over the course of the timelapse (19 hours).

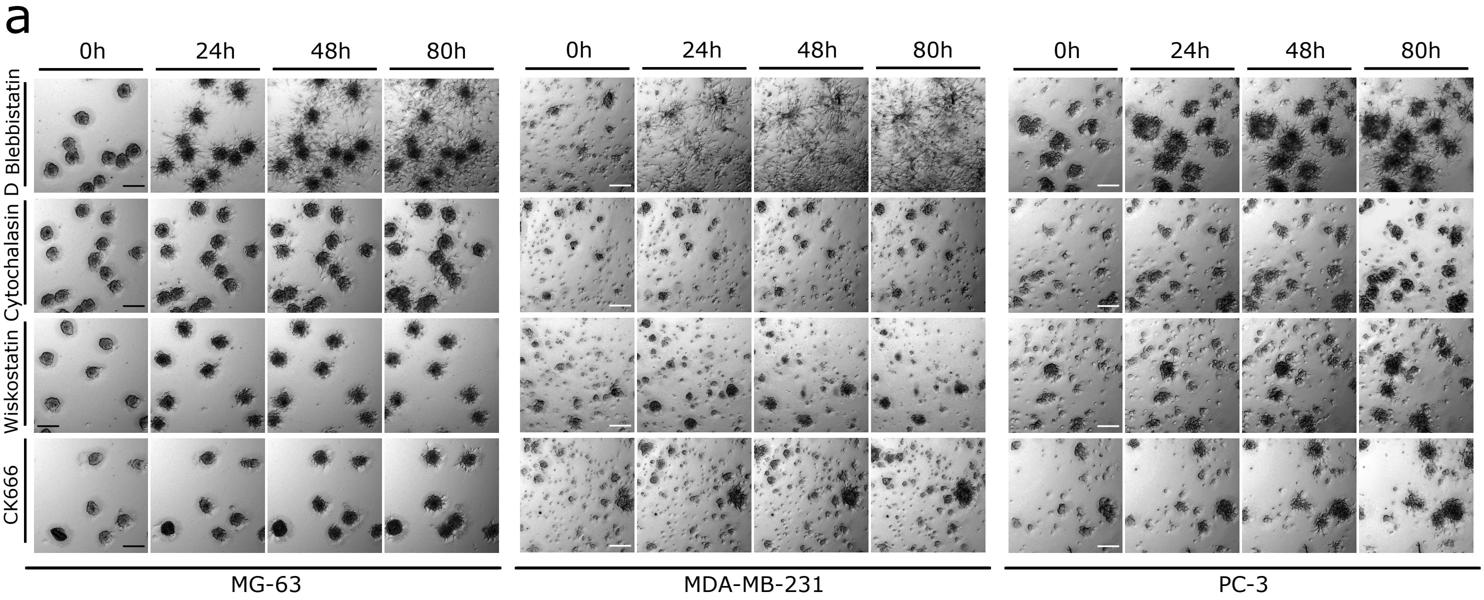


**Supplementary figure 5: (a)** Pre-formed spheroids of three representative aggregating cell lines were embedded in Matrigel and observed by means of bright-field microscopy for several days (From left to right: MG-63; MDA-MB-231 and PC-3). Pictures taken at 0, 24, 48, and 80 hours after seeding are shown. The first row of each set of images shows spheroids treated with 100 μM Blebbistatin, the second-row spheroids treated with 1 μM Cytochalasin D, the third-row spheroids treated with 10 μM Wiskostatin and the last row spheroids treated with 100 μM CK666. Seeding density: 2,5 spheroids/mm3. Scale bar: 200 μm.

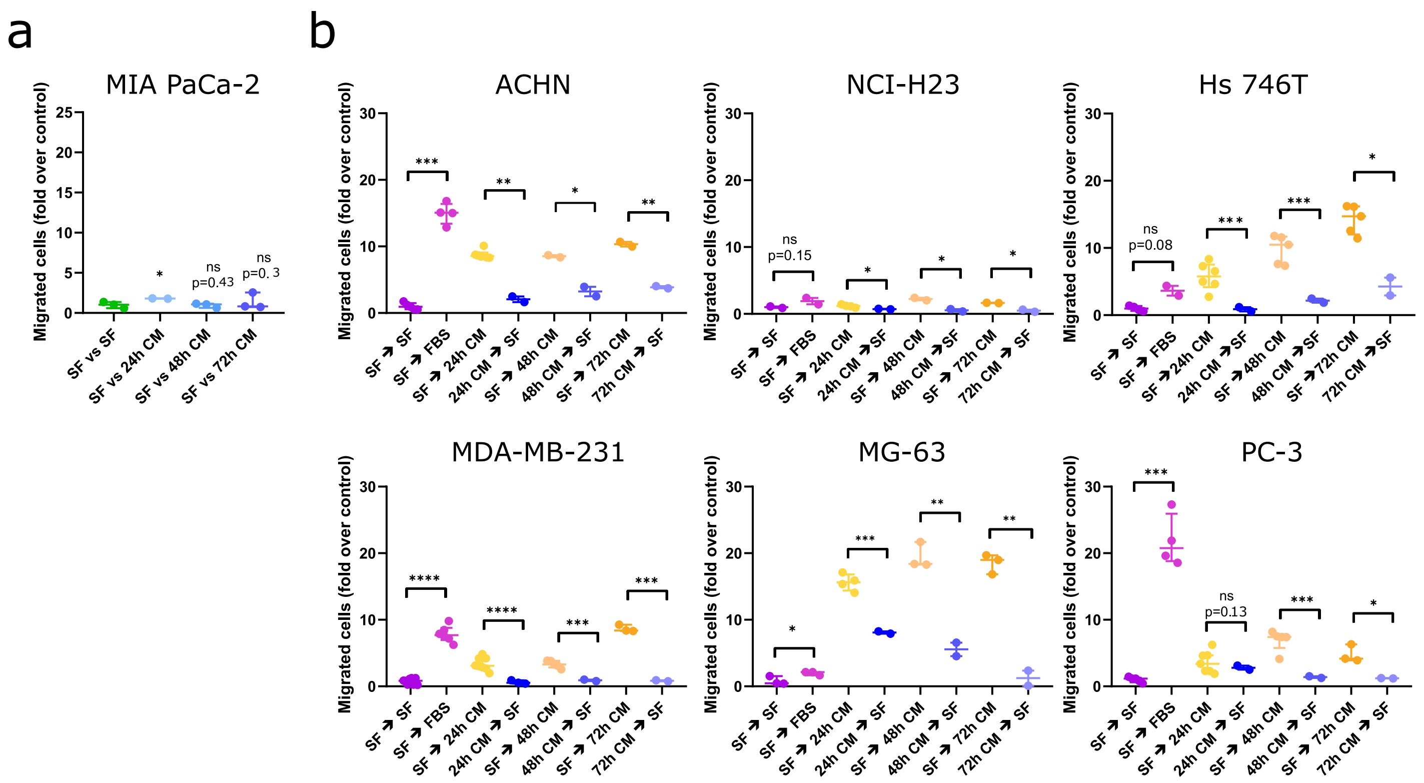


**Supplementary figure 7: (a)** Transwell assay performed with MIA PaCa-2 cells. As in Fig. 7, the plots report the migrated cells (fold over control) for conditioned media collected at different times. The data indicate that MIA PaCa-2 are not able to migrate toward their own conditioned media. Each point on the plot represents data coming from a whole membrane. Migration of cells from serum free media towards serum free media (SF - SF) was used to normalize data as a control. Data are reported as median (horizontal line) with interquartile range. Statistical significance was assessed by performing a parametric one-tailed t-test with Welch's correction (unpaired); *= P ≤ 0.05; **= P ≤ 0.01; ***= P ≤ 0.001; ****= P ≤ 0.0001. **(b)** To verify the capability of cells to migrate, we performed a control experiment by adding 1% FBS medium in the lower compartment (violet points) and use migration toward SF medium as control. Furthermore, to exclude purely chemokinetic or proliferative effects we performed the experiments by adding the conditioned medium (24, 48 and 72 hours) in the upper compartment and the serum free medium in the lower compartment. We obtained that indeed the conditioned media has a genuine effect as its presence in the upper chamber did not induce migration as in the previous conditions, or at least not to the same extent (blu-purple points versus yellow-orange points). Statistical significance between the two conditions was assessed by performing a parametric one-tailed t-test with Welch's correction (unpaired); *= P ≤ 0.05; Conditioned media are not able to attract cells. **= P ≤ 0.01; ***= P ≤ 0.001; ****= P ≤ 0.0001.

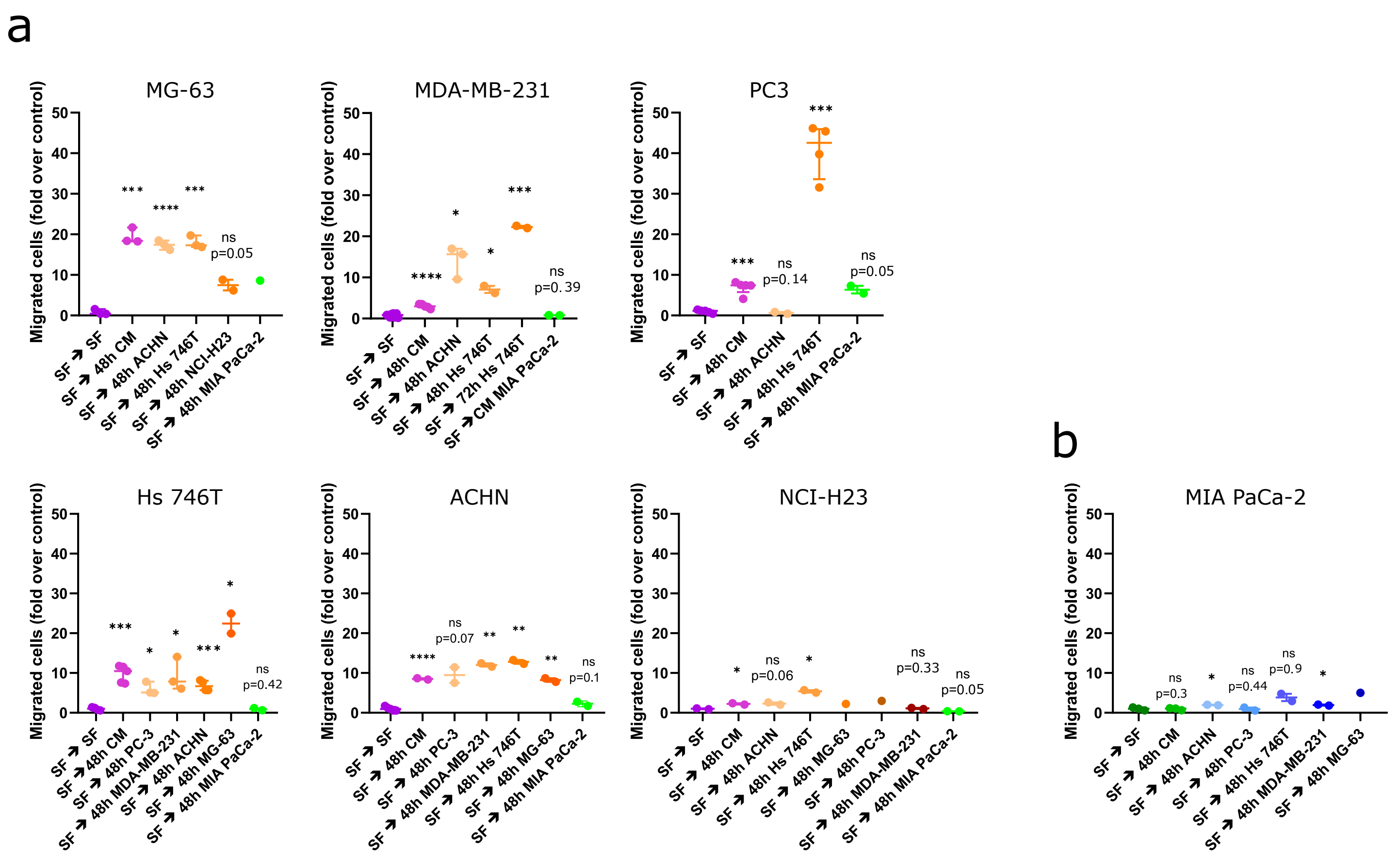


**Supplementary figure 8: (a,b)** Transwell assays performed to test whether the conditioned medium of each of the 6 aggregating cell lines, and MIA PaCa-2 as representative example of a non-aggregating cell line, act as chemoattractant for all the other cell lines. The plots report the migrated cells (fold over control) of different experiments. Each point on the plot represents a whole membrane. Data are reported as median (horizontal line) with interquartile range. Statistical significance was assessed by performing a parametric one-tailed t-test with Welch's correction (unpaired); *= P ≤ 0.05; **= P ≤ 0.01; ***= P ≤ 0.001; ****= P ≤ 0.0001.

| Cell line name | Site of origin | Culture medium |
|---|---|---|
| **786-O** | Renal cell carcinoma | RPMI |
| **A-375** | Amelanotic melanoma | DMEM |
| **A-431** | Skin squamous cell carcinoma | DMEM |
| **A-498** | Renal cell carcinoma | RPMI |
| **A549** | Lung adenocarcinoma | RPMI |
| **ACHN** | Papillary renal cell carcinoma; derived from metastatic site: pleural effusion | RPMI |
| **BxPC-3** | Pancreatic ductal adenocarcinoma | RPMI |
| **Capan-1** | Pancreatic ductal adenocarcinoma; derived from metastatic site: liver | ISCOVE 20% |
| **CFPAC-1** | Pancreatic ductal adenocarcinoma; derived from metastatic site: liver | RPMI |
| **COV-362** | High grade ovarian serous adenocarcinoma; derived from metastatic site: pleural effusion | DMEM |
| **DLD-1** | Colon adenocarcinoma | RPMI |
| **DU145** | Prostate carcinoma; derived from metastatic site: brain | RPMI |
| **GP2d** | Colon adenocarcinoma | DMEM |
| **HCT 116** | Colon adenocarcinoma | RPMI |
| **HCT-15** | Colon adenocarcinoma | RPMI |
| **HOP-62** | Lung adenocarcinoma | RPMI |
| **Hs 746T** | Gastric adenocarcinoma; derived from metastatic site: muscle; left leg | DMEM |
| **HT-29** | Colon adenocarcinoma | DMEM |
| **LNCaP** | Prostate carcinoma; derived from metastatic site: left supraclavicular lymph node | RPMI |
| **MDA-MB-231** | Breast adenocarcinoma; derived from metastatic site: pleural effusion | DMEM |
| **MG-63** | Osteosarcoma; derived from metastatic site: bone; left femur | DMEM |
| **MIA PaCa2** | Pancreatic ductal adenocarcinoma | RPMI |
| **NCI-H1299** | Lung large cell carcinoma; derived from metastatic site: lymph node | RPMI |
| **NCI-H23** | Lung adenocarcinoma | RPMI |
| **OVCAR-8** | High grade ovarian serous adenocarcinoma | RPMI |
| **PANC-1** | Pancreatic ductal adenocarcinoma | RPMI |
| **PC-3** | Prostate carcinoma; derived from metastatic site: bone | RPMI |
| **SK-OV-3** | Ovarian serous cystadenocarcinoma; derived from metastatic site: ascites | McCoy's 5A |
| **SW48** | Colon adenocarcinoma | DMEM |
| **U-2 OS** | Osteosarcoma | McCoy's |

**Table1:** list of cell lines used in this work, their tissues of origin and their culture media.

| LIGAND | RECEPTOR | REFERENCES |
|---|---|---|
| CXCL5 | CXCR1 | [1, 2] |
| | CXCR2 | [2-4] |
| CXCL6 | CXCR1 | [5, 6] |
| | CXCR2 | [2, 7] |
| CXCL1 | CXCR1, CXCR2 | [8, 9] |
| CXCL8 | CXCR1 | [5, 6] |
| | CXCR2 | [2] |
| CCL2 | CCR2 | [10] |
| CCL20 | CCR6 | [11, 12] |
| CCL20 | CCR11 | [13] |
| CCL5 | CCR1, CCR3, CCR5, CCR4 | [14-16] |
| VEGFA | FLT | [17-19] |
| | KDR | [18-20] |
| AGT | AT1, AT2 | [21] |
| DKK1 | LPR5, LPR6 | [22] |
| GDF15 | GFRAL | [23] |
| MDK | ALK | [24] |
| | NOTCH2 | [25] |
| | SDC3 | [26] |
| CCN3 | NOTCH1 | [27] |
| | ITGAV | [28] |
| | ITGA5 | [28, 29] |
| PDGFA | PDGFRA | [30] |
| GRN | TNFRSF1A, TNFRSF1B | [31, 32] |
| | DLK1 | [33, 34] |
| | EPHA2 | [35] |
| | SORT1 | [34, 36, 37] |
| IL11 | IL6ST | [38, 39] |
| | IL11RA | [40, 41] |
| IL6 | IL6R | [42, 43] |
| | IL6ST | [38, 44-46] |
| ANGPT1 | TEK | [47-49] |
| HGF | MET | [50, 51] |
| CXCL10 | CXCR3 | [52, 53] |
| IL7 | IL7R | [54] |
| XCL1 | XCR1 | [55-57] |
| PF4 | CXCR3 | [58-60] |

**Table2:** references providing evidence of receptor activity to found ligands.

**Supplementary movie legends**

**SI Movie 1:** MDA-MB-231 single cells embedded in a Matrigel and imaged once a day for 16 days after seeding. Seeding density *a posteriori*: 22,2 cells/mm$^3$. Scale bar: 200 µm.

**SI Movie 2:** PANC1 single cells embedded in a Matrigel and imaged once a day for 15 days after seeding. This cell line is reported as representative of the behaviour of non-aggregating cell lines. Seeding density *a posteriori:* 59,5 cells/mm$^3$. Scale bar: 200 µm.

**SI Movie 3:** Pre-formed spheroids of MDA-MB-231 embedded in Matrigel and imaged every 2 hours for 4 days. Seeding density: 2,5 spheroids/mm$^3$. Scale bar: 200 µm.

**SI Movie 4:** Pre-formed spheroids of MDA-MB-231 embedded in Matrigel and treated with 0.75 µg/ml mitomycin. Seeding density: 2,5 spheroids/mm$^3$. Scale bar: 200 µm.

**SI Movie 5:** Pre-formed spheroids of PC-3 embedded in Matrigel and treated with 0.75 µg/ml mitomycin. Seeding density: 2,5 spheroids/mm$^3$. Scale bar: 200 µm.

**SI Movie 6:** Pre-formed spheroids of MG-63 embedded in Matrigel and treated with 0.75 µg/ml mitomycin. Seeding density: 2,5 spheroids/mm$^3$. Scale bar: 200 µm.

**SI Movie 7:** Pre-formed spheroids of MG-63 cells transduced with LifeAct-GFP (green) or LifeAct-Ruby (red) expressing lentiviral vectors, embedded in Matrigel and imaged every hour for 3 days. Scale bar: 100 µm.

**SI Movie 8:** Pre-formed spheroids of LifeAct-Ruby expressing MDA-MB-231 embedded in Matrigel and imaged every 30 minutes for two days. Scale bar: 100 µm.

**SI Movie 9:** Pre-formed spheroids of MDA-MB-231 embedded in Matrigel and imaged every 45 minutes for 4 days. Seeding density: 2,5 spheroids/mm$^3$. Scale bar: 200 µm.

**SI Movie 10:** Pre-formed spheroids of MDA-MB-231 embedded in Matrigel, treated with 1 µm Latrunculin A and imaged every 45 minutes for 4 days. Seeding density: 2,5 spheroids/mm$^3$. Scale bar: 200 µm.

**SI Movie 11:** Pre-formed spheroids of MDA-MB-231 embedded in Matrigel, treated with 100 µm Blebbistatin and imaged every 45 minutes for 4 days. Seeding density: 2,5 spheroids/mm$^3$. Scale bar: 200 µm.

**SI Movie 12:** Pre-formed spheroids of MDA-MB-231 embedded in Matrigel, treated with 20 µM Y-27632 and imaged every 45 minutes for 4 days. Seeding density: 2,5 spheroids/mm$^3$. Scale bar: 200 µm.

**SI Movie 13:** Pre-formed spheroids of MDA-MB-231 embedded in Matrigel, treated with 10 µM Wiskostatin and imaged every 45 minutes for 4 days. Seeding density: 2,5 spheroids/mm$^3$. Scale bar: 200 µm.

**SI Movie 14:** Pre-formed spheroids of MDA-MB-231 embedded in Matrigel, treated with 10o µM CK666 and imaged every 45 minutes for 4 days. Seeding density: 2,5 spheroids/mm$^3$. Scale bar: 200 µm.

**SI Movie 15:** Pre-formed spheroids of MDA-MB-231 embedded in Matrigel, treated with 1 µM Cytochalasin D and imaged every 45 minutes for 4 days. Seeding density: 2,5 spheroids/mm$^3$. Scale bar: 200 µm.

**SI Movie 16:** Pre-formed spheroids of MDA-MB-231 embedded in Matrigel, treated with BEZ235 100 nM and imaged every 45 minutes for 4 days. Seeding density: 2,5 spheroids/mm$^3$. Scale bar: 200 µm.

**SI Movie 17:** Pre-formed spheroids of MDA-MB-231 embedded in Matrigel, treated with BYL719 3 µM and imaged every 45 minutes for 4 days. Seeding density: 2,5 spheroids/mm$^3$. Scale bar: 200 µm.

**SI Movie 18:** Pre-formed spheroids of MDA-MB-231 embedded in Matrigel, treated with AZD-6244 0,5 µM and imaged every 2 hours for 4 days. Seeding density: 2,5 spheroids/mm$^3$. Scale bar: 200 µm.

**SI Movie 19:** Pre-formed spheroids of MDA-MB-231 embedded in Matrigel, treated with MK-2206 5 µM and imaged every 45 minutes for 4 days. Seeding density: 2,5 spheroids/$mm^3$. Scale bar: 200 µm.

**SI Movie 20:** Pre-formed spheroids of H2B-RFP MDA-MB-231 (red) and H2B-GFP PC-3 (green) seeded in Matrigel. Spheroids were imaged every 2 hours for 4 days. Scale bar: 200 µm.

**SI Movie 21:** Pre-formed spheroids of H2B-RFP MDA-MB-231 (red) and H2B-GFP MG-63 (green) cell lines seeded in Matrigel. Spheroids were imaged every 2 hours for 4 days. Scale bar: 200 µm.

**SI Movie 22:** Pre-formed spheroids of H2B-RFP PC-3 (red) and H2B-GFP MG-63 (green) cell lines seeded in Matrigel. Spheroids were imaged every 2 hours for 4 days. Scale bar: 200 µm.

**SI Movie 23:** Pre-formed spheroids of LifeAct-Ruby MDA-MB-231 (red) and LifeAct-GFP PC-3 (green) cell lines in Matrigel. Spheroids were imaged every hour for 2 days. Scale bar: 100 µm. Scale bar: 100 µm.
